# TET2 loss promotes premalignant survival and clonal selection in MYC-driven B cell lymphoma

**DOI:** 10.64898/2026.03.20.712678

**Authors:** Sarah Spoeck, Nadine Kinz, Irene Rigato, Katharina Hoppe, Julia Heppke, Paul Y. Petermann, Johannes G. Weiss, Miriam Erlacher, Michael Schubert, Joel S. Riley, Andreas Villunger, Francesca Finotello, Verena Labi

## Abstract

The DNA demethylase ten-eleven translocation enzyme 2 (TET2) is frequently inactivated in hematologic malignancies, yet how its loss shapes oncogene-driven transformation remains unclear. Using a mouse model in which MYC is overexpressed in the B cell lineage, driving aggressive B cell lymphoma, we show that *Tet2* loss increases lymphoma penetrance and biases disease toward an IgM⁺ immunophenotype. Established lymphomas are broadly similar across genotypes, suggesting that *Tet2* loss exerts much of its effects before lymphoma onset. Accordingly, *Tet2* loss expands a premalignant IgM⁺ B cell subset with reduced apoptotic sensitivity and an increased frequency of BCL2⁺BIM^hi^ cells. Consistently, *Tet2* deficient IgM⁺ B cells persist better in *in vitro* cultures, show increased clonogenic survival, and exhibit clonal skewing. These findings support a model in which *Tet2* loss heightens MYC-driven lymphoma penetrance by promoting the survival and selection of premalignant B cells under apoptotic stress.

## INTRODUCTION

Hematopoiesis, the lifelong production of blood cells, depends on continuous epigenetic remodeling, in which the ten-eleven translocation enzyme 2 (TET2) plays a central role. TET2 is an iron- and α-ketoglutarate (αKG)-dependent dioxygenase that regulates gene expression by catalyzing the stepwise oxidation of 5-methylcytosine (5mC) to 5-hydroxymethylcytosine (5hmC) and further oxidized derivatives, thereby initiating DNA demethylation and restoring unmethylated cytosine^1–6^. In a cell type-specific manner, TET2 is preferentially recruited to enhancers, but can also act at promoters and gene bodies, where it helps maintain chromatin accessibility and lineage-appropriate gene expression programs^7–11^. Through these activities, TET2 is critical for proper hematopoietic differentiation, lineage commitment, and immune homeostasis, and its perturbation can have broad consequences^12–16^.

TET2 is a well-established tumor suppressor in hematopoietic cells. *Tet2*-deficient mouse models show enhanced hematopoietic stem and progenitor cell (HSPC) self-renewal, myeloid bias, altered B and T cell homeostasis, and predisposition to myeloid and lymphoid malignancies^12–15,17^. These phenotypes are commonly more pronounced when *Tet2* loss is combined with *Tet3* loss, consistent with partial functional redundancy between the two enzymes^5,8,18,19^. In B cells, TET2 and TET3 help maintain B-lineage gene regulatory programs controlled by core transcription factors (TFs) such as EBF1 and PAX5^8,11,20^. Accordingly, loss of TET activity in mouse models disrupts stage-specific enhancer demethylation, impairs B cell differentiation and function, including germinal center responses and antibody output, and predisposes to B cell malignancies^8,11,21–24^. In myeloid cells, TET deficiency causes hypermethylation of lineage-specific enhancers and of binding sites for key transcription factors such as PU.1, RUNX1 and CEBPA, thereby promoting myeloid bias, a proinflammatory state, and myeloid transformation in mice^19,25–27^.

In humans, pathogenic alterations of *TET* genes most commonly affect *TET2*, whereas they are substantially less frequent in *TET3*^5^. Rare germline *TET2* variants have been associated with immune dysregulation and predisposition to hematologic malignancy, including childhood B and T cell lymphoma^28–30^. Far more commonly, *TET2* loss-of-function (LOF) mutations arise in hematopoietic cells, often as early events in stem and progenitor compartments. They rank among the most frequent genetic alterations in human hematologic malignancies, including acute myeloid leukemia (AML), chronic myelomonocytic leukemia (CMML), myelodysplastic syndromes (MDS), myeloproliferative neoplasms (MPN), and T and B cell lymphomas^31–35^. Consistent with a role as a tumor suppressor, *TET2* LOF mutations are the second most common lesions in clonal hematopoiesis (CH), a preclinical condition in which mutant HSPC clones expand with age. Here, inflammatory cues contribute to the selective expansion of *TET2*-deficient HSPCs, and may facilitate malignant transformation^27,36,37^.

Across mouse models of myeloid malignancy, *Tet2* deficiency recurrently cooperates with oncogenes such as FLT3-ITD, JAK2^V617F^, or RAS to shorten disease latency or increase penetrance^31,38–42^. Additional cooperating events include epigenetic modifiers such as DNMT3A or ASXL1, as well as loss of the tumor suppressor TP53^43–46^. To date, only a few studies have addressed cooperation of *Tet2* loss with additional oncogenic lesions in B cells, notably BCL6 and TCL1A, where it accelerates lymphomagenesis^16,47^. However, whether *Tet2* loss similarly cooperates with other oncogenic drivers to promote B cell transformation remains unknown, despite recurrent *TET2* LOF mutations in human B cell malignancies. This gap is particularly notable given the central role of oncogenic MYC deregulation in aggressive human B cell lymphomas, including Burkitt lymphoma (BL) and diffuse large B-cell lymphoma (DLBCL)^48,49^. Although physiologically required for hematopoietic progenitor function and early B cell development, enforced MYC expression imposes strong proliferative pressure while sensitizing cells to apoptosis, a constraint that must be overcome during transformation^50–54^. This raises the question of how *Tet2* loss shapes premalignant and malignant B cell states during MYC-driven lymphomagenesis *in vivo*.

To investigate this, we used the *EμMyc* mouse model, in which enforced MYC expression in the B cell lineage drives aggressive lymphoma^55,56^. TET2 deficiency increased MYC-driven B cell lymphoma penetrance and shifted the tumor spectrum toward IgM⁺ disease. Although established lymphomas were broadly similar across genotypes, premalignant *EμMyc Tet2^−/−^*mice showed a partial block in peripheral B cell maturation, reflected by accumulation of IgM⁺ immature-like B cells that are largely CD21*^−^* and CD23*^−^*. Within this compartment, *Tet2* loss enriched a BCL2⁺BIM^hi^ subpopulation associated with enhanced *in vitro* survival, increased functional BCL2-dependence, greater colony-forming capacity, and increased clonal skewing. Together, these findings identify *Tet2* loss as a cooperating lesion that facilitates MYC-driven lymphomagenesis by restraining B cell maturation, buffering apoptotic stress, and promoting early clonal skewing.

## RESULTS

### *Tet2* loss increases B cell lymphoma penetrance in *EμMyc* mice

To test how TET2 LOF influences MYC-driven lymphomagenesis, we monitored mouse cohorts with germline *Tet2* deletion^13^ with or without the B cell-specific *EμMyc* transgene^55,56^. In the absence of MYC overexpression, *wildtype*, *Tet2^+/−^*, and *Tet2^−/−^* animals remained free of overt disease over the observation period of 300 days (Fig. 1A). As expected, *EμMyc* mice developed aggressive B cell lymphoma with a median overall survival of 127 days, and 26.8% of animals remained tumor-free until day 300 (Fig. 1A). *Tet2* loss reduced overall survival on the *EμMyc* background, with 122 days for *EμMyc Tet2^+/−^* mice and 103 days for *EμMyc Tet2^−/−^* mice (Fig. 1A). Notably, all *EμMyc Tet2^−/−^*mice developed tumors, whereas long-term tumor-free survivors persisted in the *EμMyc* and *EμMyc Tet2^+/−^* cohorts. Disease latency of the animals that ultimately succumbed was equal across genotypes (Fig. 1B*; EμMyc* 105 days; *EμMyc Tet2^+/−^* 108 days; *EμMyc Tet2^−/−^* 103 days), indicating that *Tet2* loss primarily increases lymphoma penetrance upon oncogenic MYC expression.

**Figure 1:**
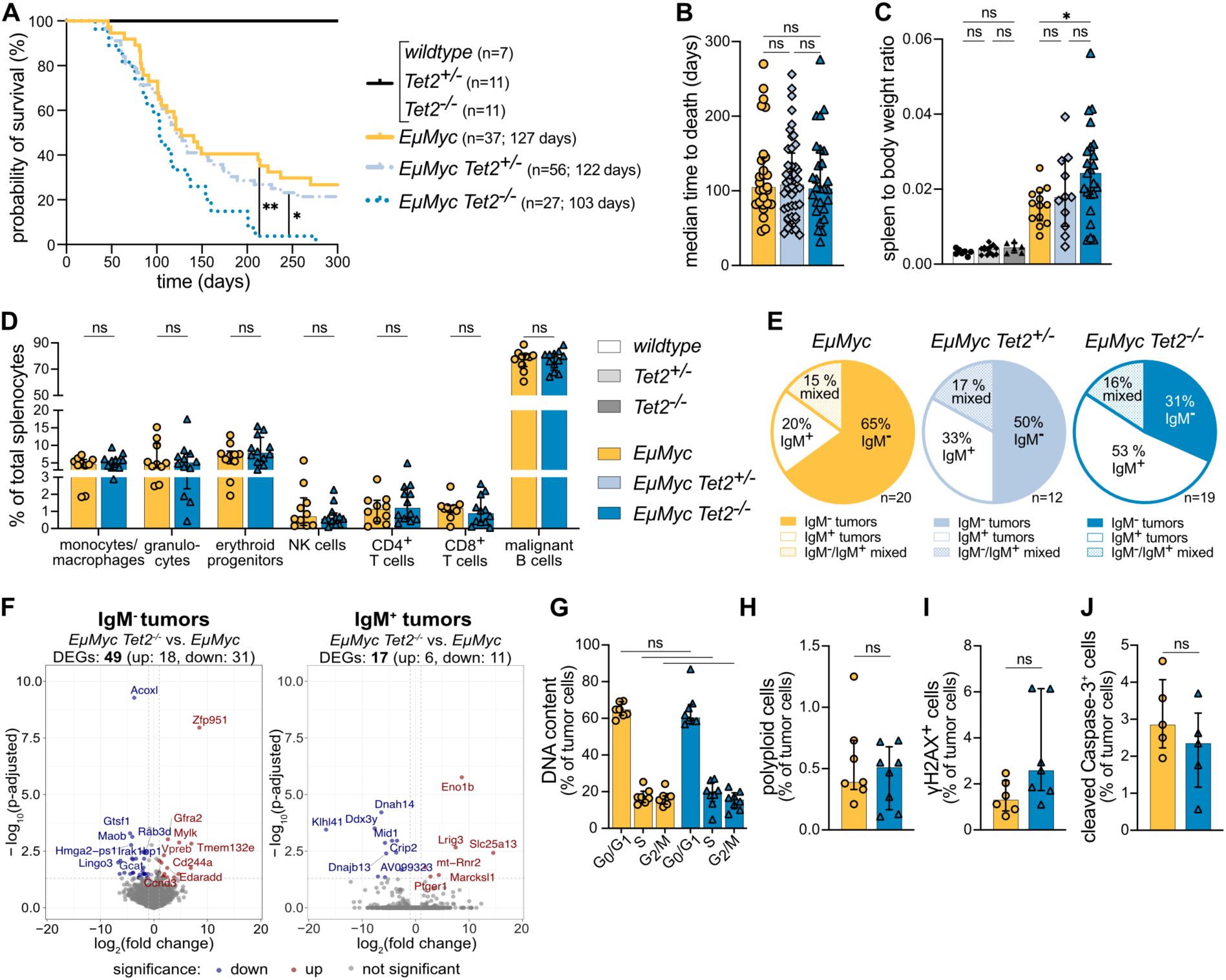
*Tet2* loss increases B cell lymphoma penetrance in *EμMyc* mice. Kaplan-Meier survival analysis displaying **(A)** overall survival probability, and **(B)** median time to death in days for all malignant mice (*EµMyc*: n=27, *EµMyc Tet2^+/−^*: n=44, *EµMyc Tet2^−/−^*: n=27). **(C)** Spleen to body weight ratio in non-malignant (280-320 days) mice (*wildtype*: n=7*, Tet2^+/−^*: n=13*, Tet2^−/−^*: n=6) and malignant mice, excluding animals without overt disease at 300 days (*EµMyc*: n=13, *EµMyc Tet2^+/−^*: n=11, *EµMyc Tet2^−/−^*: n=23). **(D)** Splenic immune cell composition assessed by flow cytometry: monocytes/macrophages (CD11b^+^Gr1^−^), granulocytes (CD11b^+^Gr1^+^), erythroid progenitors (nucleated Ter119^+^), NK cells (TCRβ^−^NK1.1^+^), CD4^+^ T cells (TCRβ^+^CD4^+^), CD8^+^ T cells (TCRβ^+^CD8^+^), and malignant B cells (B220^+^CD19^+^) (*EµMyc*: n=10, *EµMyc Tet2^−/−^*: n=12). **(E)** Lymphoma immunophenotype in the spleen. Mixed tumors were defined as tumors where neither IgM^−^ nor IgM^+^ cells constituted >80% of the total tumor population. **(F)** Volcano plots displaying RNA-seq-derived transcriptional profiles of FACS-sorted IgM^−^ tumor (left; B220^+^CD19^+^IgM^−^IgD^−^) and IgM^+^ tumor (right; B220^+^CD19^+^IgM^+^IgD^−^) cells. Comparison between *EµMyc Tet2^−/−^* and *EµMyc* (IgM^−^ tumors: n=4 vs. n=5, IgM^+^ tumors: n=4 vs. n=3) mice was performed separately for each cell type. Significance was defined as adjusted p-value<0.05 and absolute log_2_(fold change)>1. Downregulated genes in *EµMyc Tet2^−/−^* tumors are shown in blue; upregulated genes are shown in red. **(G)** DNA content and **(H)** polyploid cell fraction of splenic lymphoma cells, assessed by TO-PRO-3 staining via flow cytometry (*EµMyc*: n=7, *EµMyc Tet2^−/−^*: n=8). Flow cytometry assessment of **(I)** DNA double strand breaks by γH2AX staining (*EµMyc*: n=6, *EµMyc Tet2^−/−^*: n=7) and **(J)** apoptosis by cleaved Caspase-3 staining (*EµMyc*: n=5, *EµMyc Tet2^−/−^*: n=5) within splenic lymphoma cells. Bar graphs show median with interquartile range. Statistical significance was determined using (A) Mantel-Cox test, (B, C) one-way ANOVA, (D, G) Mann-Whitney test, or (H, I, J) unpaired t-test, depending on normality (Shapiro-Wilk test) with Holm-Šidák correction for multiple comparisons. ns = not significant, *p<0.05, **p<0.005.

To assess tumor burden at two major sites of disease involvement in the *EμMyc* model, we analyzed spleens and bone marrow. As expected, spleen to body weight ratio was elevated in tumor-bearing *EμMyc* mice compared with non-transgenic controls, however, *Tet2* loss increased spleen weight further (Fig. 1C), and total splenocyte and bone marrow cell numbers showed a similar trend (Fig. S1A). Flow cytometric profiling of spleen and bone marrow composition revealed comparable frequencies of major immune lineages and malignant B cells between *EμMyc* and *EμMyc Tet2^−/−^* mice (Fig. 1D and S1B), suggesting that germline *Tet2* loss does not induce major shifts in overall immune composition in established disease.

We next classified tumors by surface B cell receptor (BCR) expression of the immunoglobulin M (IgM) isotype, which is commonly used to stratify *EμMyc* lymphomas by cell-of-origin^55,57^. IgM⁻ tumors are typically linked to transformation within a proliferating, developmentally arrested compartment of IgM⁻ progenitor B cells^55,57^. IgM⁺ tumor cells are commonly IgD⁻ and show minimal IgV (Immunoglobulin variable region) somatic hypermutation with low activation-induced cytidine deaminase (AID) expression, while recombinant activating gene (RAG) activity has been reported, supporting an immature-like, proliferating pre-germinal center state as cell-of-origin^58,59^. A majority of *EμMyc* mice presented IgM⁻ tumors, while a smaller proportion developed IgM⁺ or IgM⁻/IgM⁺ mixed disease (Fig. 1E and S1C). In contrast, *EμMyc Tet2^−/−^* mice predominantly developed IgM⁺ tumors, and *EμMyc Tet2^+/−^*animals showed an intermediate phenotype (Fig. 1E and S1C). This pattern is consistent with a gene-dosage effect, as also reported in other hematopoietic mouse models^5,60^. Together, these data identify TET2 as a suppressor of MYC-driven transformation that appears particularly relevant within the BCR-expressing compartment.

To assess whether *Tet2* loss is associated with persistent transcriptional differences in established disease, we performed bulk RNA-seq on fluorescence-activated cell sorting (FACS)-purified IgM⁻ and IgM⁺ tumor cells. In both immunophenotypic subsets, comparing *EμMyc Tet2^−/−^* with *EμMyc* tumors revealed only a small number of significantly differentially expressed genes (DEGs) (Fig. 1F and Table S1). Hallmark gene set analyses likewise showed no major genotype-associated changes (Fig. S1D). These data argue against widespread transcriptional remodeling in established *Tet2*-deficient tumors.

Given prior work linking *Tet2* or combined *Tet2/Tet3* loss to altered expression of DNA repair genes and increased DNA damage in hematologic malignancies^17,19,61,62^, we examined proliferative state and genomic integrity in established lymphomas *ex vivo*. DNA content profiling revealed similar cell cycle distribution across G_0_/G_1_, S, and G_2_/M phases between *EμMyc* and *EμMyc Tet2^−/−^* tumors (Fig. 1G and Fig. S1E). Polyploidy was rare but comparable between genotypes (Fig. 1H and Fig. S1F). We also inferred copy-number profiles from our bulk RNA-seq data, which revealed heterogeneous aneuploidy patterns across tumors without a consistent genotype-associated signature (Fig. S1G). Accordingly, Hallmark gene set analyses did not indicate genotype-dependent changes in DNA repair programs (Fig. S1D and Table S1), and γH2AX staining did not demonstrate a significant increase in DNA double-strand breaks in *Tet2*-deficient tumor cells compared with controls (Fig. 1I and Fig. S1H). Finally, fractions of tumor cells with cleaved Caspase-3 were comparable between genotypes (Fig. 1J and Fig. S1I), indicating that *Tet2* loss does not substantially alter baseline apoptosis in established lymphomas.

Overall, *Tet2* loss increases *EμMyc* lymphoma penetrance and raises the fraction of IgM⁺ tumors, while established lymphomas are largely similar across genotypes.

### *Tet2* loss enriches for IgM⁺IgD⁻ immature-like B cells in premalignant *EμMyc* mice

To determine whether the premalignant phase already shows changes that could underlie the later enrichment of IgM⁺ tumors, we analyzed animals at day 50, when most mice are still without overt disease. *EμMyc Tet2^−/−^* mice exhibited a mild increase in spleen to body weight ratio and splenocyte number compared with *EμMyc* controls (Fig. 2A, B). Flow cytometric profiling of the spleen did not reveal major genotype dependent shifts in the relative abundance of major immune cell subsets in the *EμMyc* context, including premalignant B cells (Fig. 2C). In non-*EμMyc* littermate controls, immune subset frequencies were likewise largely unchanged, aside from a modest increase in monocytes/macrophages in *Tet2^−/−^* controls (Fig. S2A), consistent with prior reports^12,14^. Interestingly, however, specific to the *EμMyc* context *Tet2* loss skewed peripheral B cell maturation in the spleen (Fig. 2D). Thus, *EμMyc Tet2^−/−^*mice showed an increased fraction of IgM⁺IgD⁻ immature-like B cells and a marked reduction of IgM⁺IgD⁺ mature B cells, while changes in the IgM⁻ progenitor B cell fraction were modest. In the bone marrow, B cells were dominated by IgM⁻ progenitors, with only minor differences in the IgM⁺IgD⁻ immature-like fraction and the expected underrepresentation of IgM⁺IgD⁺ mature B cells in the presence of oncogenic MYC (Fig. S2B). This fits normal B cell biology, where IgM⁺IgD⁻ immature B cells accumulate in the spleen during peripheral maturation, a bias that is also evident in *EμMyc* mice and further enhanced by *Tet2* loss.

**Figure 2:**
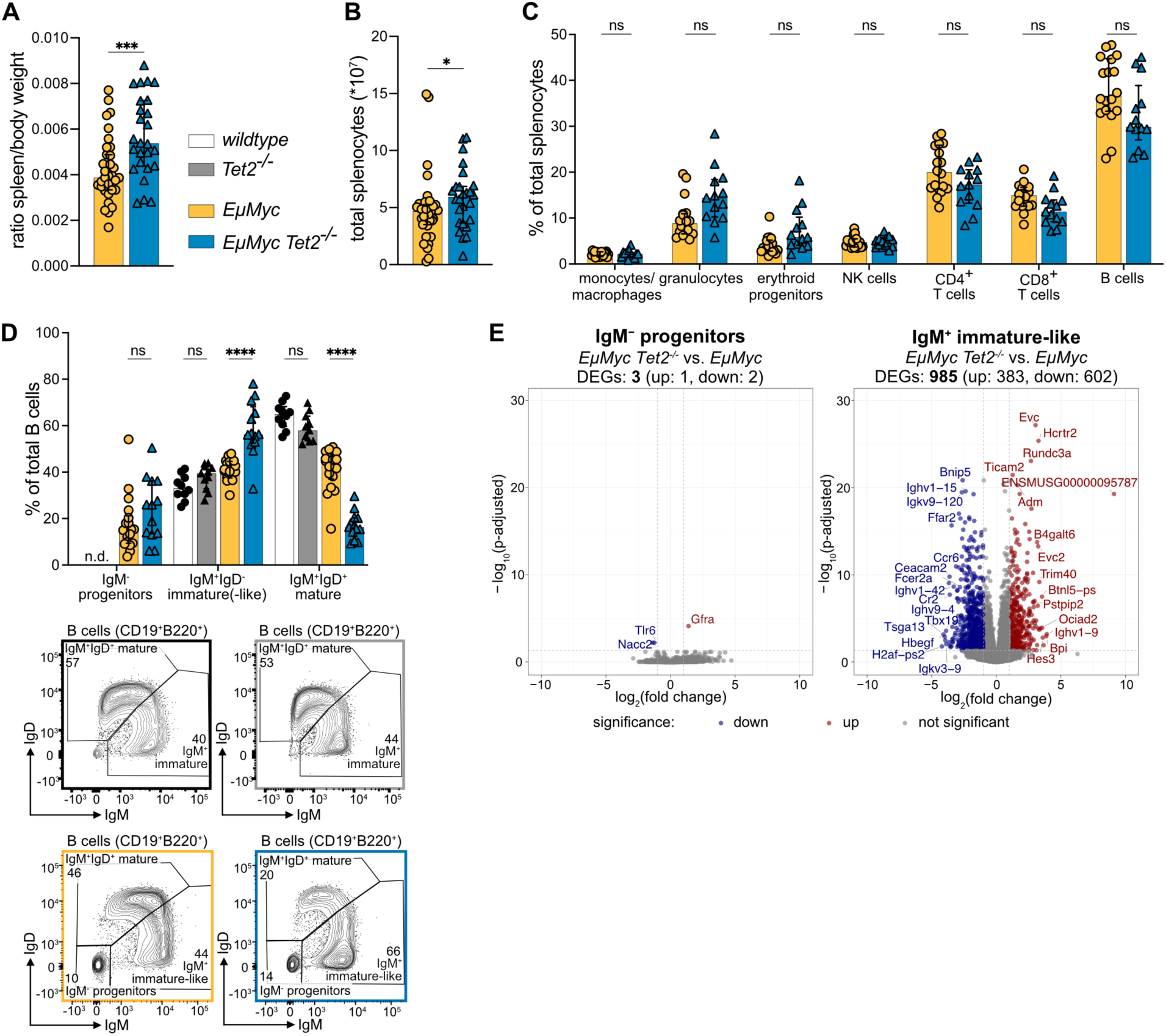
*Tet2* loss enriches for IgM⁺IgD⁻ immature-like B cells in premalignant *EμMyc* mice. Analysis of splenocytes from premalignant (day 50) *EµMyc* and *EµMyc Tet2^−/−^* mice. **(A)** Spleen to body weight ratio for each mouse and **(B)** absolute splenocyte counts (*10^7^) (*EµMyc*: n=35, *EµMyc Tet2^−/−^*: n=26). **(C)** Splenic immune cell composition assessed by flow cytometry: monocytes/macrophages (CD11b^+^Gr1^−^), granulocytes (CD11b^+^Gr1^+^), erythroid progenitors (nucleated Ter119^+^), NK cells (TCRβ^−^NK1.1^+^), CD4^+^ T cells (TCRβ^+^CD4^+^), CD8^+^ T cells (TCRβ^+^CD8^+^), and B cells (B220^+^CD19^+^) (*EµMyc*: n=18, *EµMyc Tet2^−/−^*: n=13). **(D)** Splenic B cell subsets were assessed via flow cytometry: IgM^−^ progenitors (B220^+^CD19^+^IgM^−^IgD^−^), IgM^+^IgD^−^ immature-(like) (B220^+^CD19^+^IgM^+^IgD^−^), and IgM^+^IgD^+^ mature (B220^+^CD19^+^IgM^+^IgD^+^) B cells. The upper panel summarizes all data (*wildtype*: n=10, *Tet2^−/−^*: n=11, *EµMyc*: n=18, *EµMyc Tet2^−/−^*: n=13), the lower panel shows representative dot blots for the IgM/IgD gate. **(E)** Volcano plots display RNA-seq-derived transcriptional profiles of premalignant FACS-sorted splenic IgM^−^ progenitors (left; B220^+^CD19^+^IgM^−^IgD^−^) and IgM^+^ immature(-like) (right; B220^+^CD19^+^IgM^+^IgD^−^) B cells. Comparisons between *EµMyc Tet2^−/−^* (n=5) and *EµMyc* (n=6) mice were performed separately for each cell type. Axis ranges were kept identical across volcano plots to allow direct comparison. Significance was defined as adjusted p-value<0.05 and absolute log_2_(fold change)>1. Downregulated genes in *EµMyc Tet2^−/−^*subsets are shown in blue; upregulated genes are shown in red. Bar plots show median with interquartile range. Statistical significance was determined using (A) unpaired t-test or (B, C) Mann-Whitney test and (D) two-way ANOVA, with Holm-Šidák correction for multiple comparisons. Normality was assessed using the Shapiro-Wilk test. n.d. = not detected, ns = not significant, *p<0.05, **p<0.005, ***p<0.0005.

To gain molecular insight into this phenotype, we FACS-sorted IgM⁻ progenitors and IgM⁺ immature-like B cells from premalignant *EμMyc* and *EμMyc Tet2^−/−^* spleens and performed bulk RNA-seq. In IgM⁻ progenitors, genotype-associated differences were minimal, with only 3 DEGs (Fig. 2E and Table S2). By contrast, *Tet2* loss was associated with extensive transcriptional changes in IgM⁺ immature-like B cells, with 985 DEGs (Fig. 2E and Table S2). In the non-*EμMyc* context, the same analysis revealed only few DEGs in the corresponding B cell subsets (Fig. S2C and Table S3). Altogether, the skewed peripheral B cell maturation and the pronounced transcriptional phenotype in *EμMyc Tet2^−/−^*mice depend on aberrant MYC expression and are not a dominant effect of *Tet2* loss alone.

### *Tet2* loss skews IgM⁺ immature-like B cell sub-states on the *EμMyc* background

Given the extensive transcriptional changes in premalignant IgM⁺ immature-like B cells (Fig. 2E), we next asked which gene programs are most perturbed by *Tet2* loss. GO-term enrichment analysis of the DEG set highlighted biological processes linked to B cell differentiation/maturation and immune receptor/immunoglobulin programs (Fig. 3A). This is consistent with altered maturation programs within this subset. To infer TF activity from the RNA-seq profiles we performed “footprint” analysis using decoupleR^63^. This revealed a marked reduction in E2A/TCF3 (E12/E47)-associated activity in *EμMyc Tet2*^−/−^ IgM⁺ immature-like B cells (Fig. 3B), alongside increased activity scores linked to KLF4, JUNB, NR4A1, and RUNX1. Together, these shifts are consistent with reduced maturation-associated activity (TCF3/E2A) and increased stimulus-response/state-regulatory programs (KLF4, JUNB, NR4A1, RUNX1). Because no single widely adopted signature robustly captures the murine IgM⁺ immature-to-mature transition, we curated a focused B-lineage regulatory/maturation gene panel comprising core B-lineage transcription factors, canonical peripheral maturation markers, and signaling/activation nodes. Within this panel, genotype-dependent changes were restricted to a subset of genes (13 up, 14 down, 76 unchanged) (Fig. 3C), supporting a selective perturbation rather than a global collapse of the B cell maturation program.

**Figure 3:**
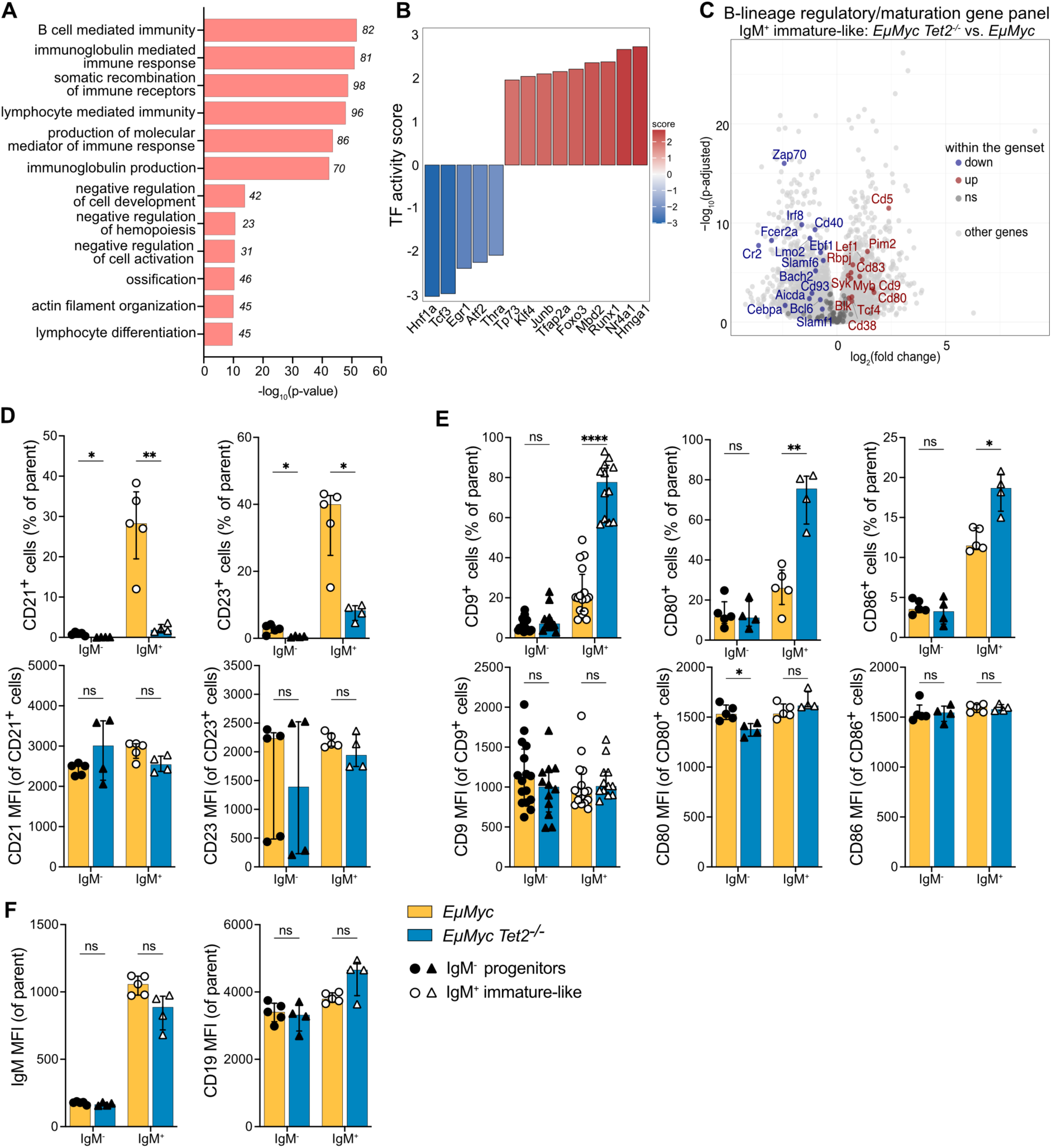
*Tet2* loss skews IgM⁺ immature-like B-cell sub-states on the *EμMyc* background. **(A)** GO-term enrichment analysis of RNA-seq data from FACS-sorted IgM^+^ immature-like B cells from the comparison of premalignant *EµMyc Tet2^−/−^* (n=5) versus *EµMyc* (n=6) mice. DEGs (adjusted p-value<0.05 and absolute log_2_(fold change)>1) were subjected to GO-term biological process analysis in RStudio. The bar graph depicts the top enriched biological processes ranked by adjusted p-value, with the number of DEGs contributing to each term indicated in italics at the end of each bar. **(B)** DecoupleR analysis on the RNA-seq experiment performed in RStudio using the CollecTRI transcription factor (TF)-target network. Bar plot displays the top 14 TFs with at least 45 targets and p-value<0.05. **(C)** Volcano plot comparing transcriptional profiles, with a specific focus on a self-defined B-lineage regulatory/maturation gene panel from the RNA-seq analysis described in (A). Significance was defined as adjusted p-value<0.05 and absolute log2(fold change)>0.5. Downregulated genes in *EµMyc Tet2^−/−^* subsets are shown in blue; upregulated genes are shown in red; B-lineage regulatory/maturation genes, which are not differentially expressed, are shown in dark grey, and all other genes in bright grey. **(D-E)** Flow cytometric validation of selected B-lineage regulatory and maturation genes in IgM^−^ progenitors (filled symbols; B220^+^CD19^+^IgM^−^IgD^−^) and IgM^+^ immature-like B cells (empty symbols; B220^+^CD19^+^IgM^+^IgD^−^). The upper panel displays the percentage of cells expressing the indicated markers, while the lower panel shows the geometric MFI within the respective marker-positive gate. **(F)** MFI of surface IgM and CD19 within IgM^−^ progenitors (filled symbols; B220^+^CD19^+^IgM^−^IgD^−^) and IgM^+^ immature-like B cells (empty symbols; B220^+^CD19^+^IgM^+^IgD^−^). Bar plots show median with interquartile range. Statistical significance was assessed using unpaired t-test for %CD21^+^ and CD21 MFI, %CD23^+^, %CD80^+^, %CD86^+^, IgM MFI, CD19 MFI, or Mann-Whitney test for CD23 MFI, %CD9^+^ and CD9 MFI, CD80 MFI, CD86 MFI, with Holm-Šidák correction for multiple comparisons. Normality was evaluated using the Shapiro-Wilk test. MFI = mean fluorescence intensity, ns = not significant, *p<0.05, **p<0.005, ****p<0.0001.

We next validated cell surface markers identified as DEGs in Fig. 3C via flow cytometry in IgM⁺ immature-like B cells, using the IgM⁻ progenitor compartment as an internal control. At the protein level, the most prominent changes were reduced frequencies of CD21/*Cr2*⁺ and of CD23/*Fcer2a*⁺ cells within the IgM^+^ immature-like compartment in *EμMyc Tet2*^−/−^ mice (Fig. 3D), two markers typically upregulated during splenic peripheral B cell maturation toward the IgM^+^IgD^+^ mature stage^64,65^. CD21 and CD23 mean fluorescence intensities (MFIs) within their respective marker-positive IgM⁺ gates were comparable between genotypes (Fig. 3D). *Aicda* and *Bcl6* expression was low in *EμMyc* IgM⁺ immature-like B cells and further decreased upon *Tet2* loss, consistent with a pre-germinal center state (Fig. S3B, C and Table S2). We then extended this analysis to additional markers identified as DEGs in Fig. 3C to further resolve heterogeneity within the IgM⁺ immature-like compartment beyond CD21 and CD23. *Tet2* loss primarily enhanced the frequencies of CD9⁺, CD80⁺ and CD86⁺ subsets (linked to co-stimulatory and interaction programs), with modest changes in the CD5⁺, CD38⁺, CD40⁺ and CD44⁺ subsets (associated with broader signaling and interaction programs) (Fig. 3E and Fig. S3A). Again, MFIs within the respective marker-positive gates were largely comparable between genotypes (Fig. 3E and Fig. S3A), suggesting that *Tet2* loss mainly alters subpopulation frequencies rather than per-cell expression.

Finally, as apparent shifts along the peripheral maturation axis can be confounded by changes in core B cell identity or BCR surface abundance, we assessed CD19 and IgM in the IgM⁺ immature-like gate. Because CD19 and surface IgM MFIs were comparable between genotypes (Fig. 3F), we next asked whether downstream BCR signaling may be altered in *EμMyc Tet2*^−/−^ IgM⁺ immature-like B cells.

### *Tet2* loss selects an apoptosis-buffered IgM⁺ B cell sub-state on the *EμMyc* background

To profile BCR-associated signaling, we performed intracellular flow cytometry in *EμMyc* and *EμMyc Tet2*^−/−^ IgM⁺ immature-like splenic B cells at steady state. We observed enhanced abundance of the non-phosphorylated proteins CD79A and SYK in *EμMyc Tet2*^−/−^ cells (Fig. S4). Overall, these changes were modest and affected proximal BCR-associated proteins without a corresponding shift across the phosphorylation readouts. Interestingly, we also observed elevated phosphorylated AKT (pAKT) in *EμMyc Tet2*^−/−^ cells (Fig. S4), a readout that can reflect BCR-PI3K signaling but also signals from other growth and survival pathways^66,67^. Thus, we used Hallmark gene set enrichment analysis on the RNA-seq data from the premalignant IgM⁺ immature-like B cells to identify transcriptional programs accompanying this change. The top enriched pathway was IL-2/STAT5 signaling, with additional enrichment of IL-6/JAK/STAT3 signaling and TNFα signaling via NF-κB in *EμMyc Tet2*^−/−^ relative to *EμMyc* cells (Fig. 4A). In this premalignant setting, we interpret these signatures as a MYC-linked stress/survival response rather than overt cytokine stimulation. Among the top hits were also p53 pathway and apoptosis (Fig. 4A), in line with prior reports on MYC-driven proliferative pressure and mitochondrial apoptotic priming^68,69^. BCL2-family transcript analyses further showed increased expression of pro-apoptotic BH3-only genes (including *Bcl2l11* and *Bbc3*) alongside elevated anti-apoptotic factors (including *Bcl2* and *Bcl2a1* isoforms) in *EμMyc Tet2*^−/−^ IgM⁺ immature-like cells (Fig. 4B). Thus, *Tet2* loss is associated with co-induced pro- and anti-apoptotic BCL2-family signatures, consistent with enrichment of an apoptosis-primed yet buffered sub-state^68,70^.

**Figure 4:**
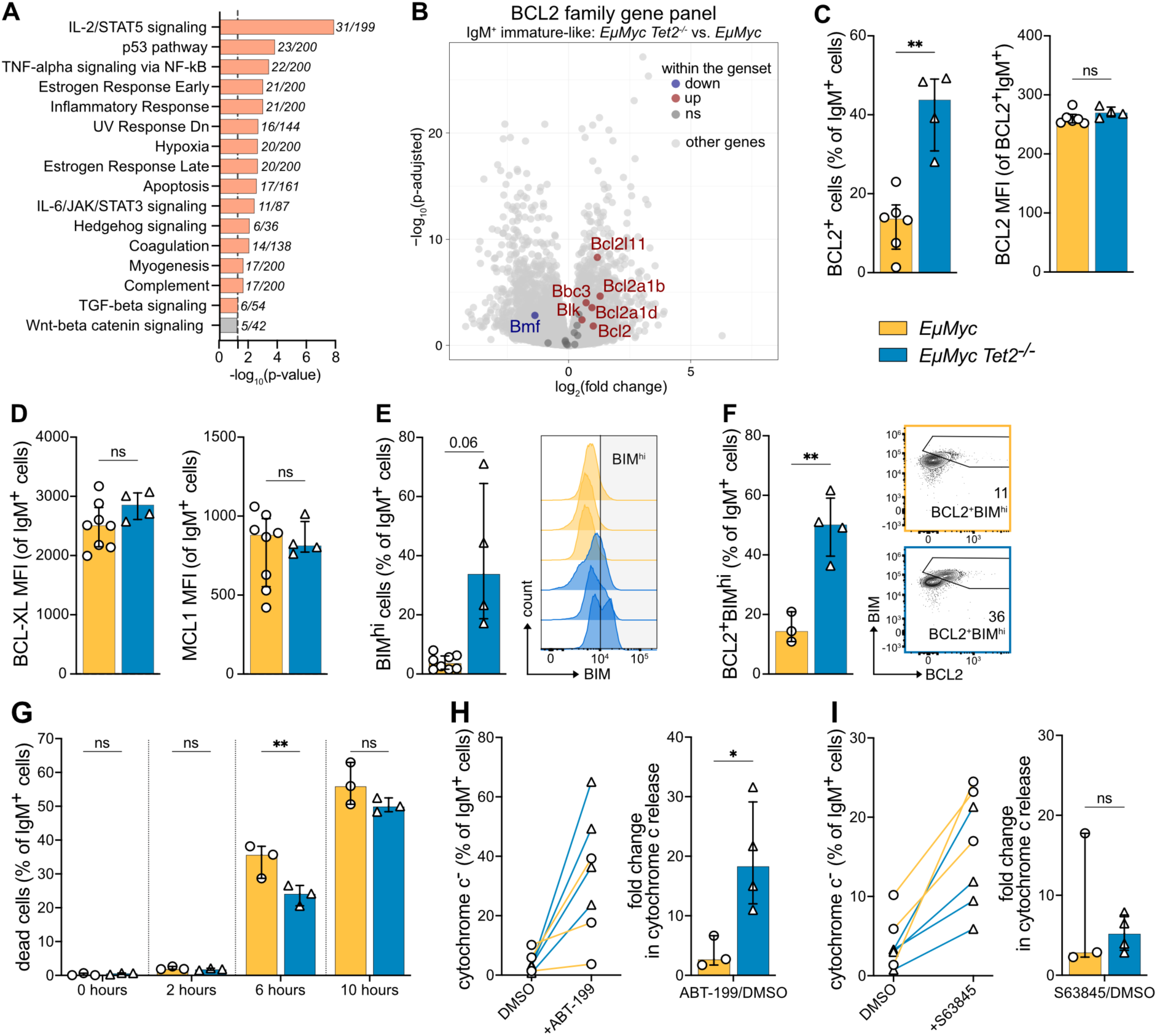
*Tet2* loss selects an apoptosis-buffered IgM⁺ B cell sub-state on the *EμMyc* background. **(A)** GO-term analysis of RNA-seq data from FACS-sorted IgM^+^ immature-like B cells from the comparison of premalignant *EµMyc Tet2^−/−^* (n=5) versus *EµMyc* (n=6) mice. DEGs (adjusted p-value<0.05 and absolute log_2_(fold change)>1) were subjected to MSigDB Hallmark 2020 in Enrichr. The bar graph depicts the top enriched pathways ranked by p-value, with the number of DEGs contributing to each term indicated in italics at the end of each bar. **(B)** Volcano plot comparing transcriptional profiles, with a specific focus on apoptotic/BCL2-family gene panel from the RNA-seq analysis described in (A). Significance was defined as adjusted p-value<0.05 and absolute log_2_(fold change)>0.5. Downregulated genes in *EµMyc Tet2^−/−^*subsets are shown in blue; upregulated genes are shown in red; apoptotic/BCL2-family genes, which are not differentially expressed in dark grey, and all other genes in bright grey. **(C)** Flow cytometric analysis determining the fraction of BCL2^+^ cells and MFI within the BCL2^+^ gate in IgM^+^ immature-like B cells (B220^+^CD19^+^IgM^+^IgD^−^) (*EµMyc*: n=6, *EµMyc Tet2^−/−^*: n=4). **(D)** MFI of BCL-XL and MFI of MCL1 in IgM^+^ immature-like B cells, quantified by flow cytometry (*EµMyc*: n=8, *EµMyc Tet2^−/−^*: n=4). **(E)** Flow cytometric analysis determining the fraction of BIM^hi^ cells and representative histograms for the BIM staining in the IgM^+^ immature-like B cell compartment (*EµMyc*: n=8, *EµMyc Tet2^−/−^*: n=4). **(F)** Flow cytometric assessment determining the fraction of BCL2^+^BIM^hi^ cells and representative dot plots of the BCL2^+^BIM^hi^ population in IgM^+^ immature-like B cells (*EµMyc*: n=3, *EµMyc Tet2^−/−^*: n=4). **(G)** Cell survival kinetics assessed *in vitro* for IgM^+^ immature-like B cells (DAPI^+^B220^+^CD19^+^IgM^+^IgD^−^) from *EµMyc* (n=3) and *EµMyc Tet2^−/−^* (n=3) mice, at 0, 2, 6, and 10 hours of culture. Assessment of mitochondrial apoptotic sensitivity via cytochrome c release of IgM^+^ immature-like B cells (ZombieDye^−^B220^+^CD19^+^IgM^+^IgD^−^cytochromec^−^) treated with **(H)** 30 µM ABT-199/Venetoclax and **(I)** 1 µM S63845. For both treatments, cytochrome c release in DMSO controls and treated samples is shown as a line plot (left) and as a bar graph (right), depicting the fold change relative to DMSO (*EµMyc*: n=3, *EµMyc Tet2^−/−^*: n=4). Bar plots show median with interquartile range. Statistical significance was assessed using unpaired t-test (A-F, H, I), or two-way ANOVA (G) with Holm-Šidák correction for multiple comparisons. Normality was evaluated using the Shapiro-Wilk test. MFI = mean fluorescence intensity, ns = not significant, *p<0.05, **p<0.005.

At the protein level, intracellular flow cytometry revealed an increased fraction of BCL2⁺ IgM⁺ immature-like cells in *EμMyc Tet2*^−/−^ compared with *EμMyc* mice, while the BCL2 MFI within the BCL2⁺ gate was comparable between genotypes (Fig. 4C). MCL1 and BCL-XL levels were unchanged (Fig. 4D). Pro-apoptotic BIM, the product of the *Bcl2l11* gene, was detected across essentially all cells, but a distinct BIM^hi^ subpopulation was evident predominantly in *Tet2* deficient *EμMyc* cells (Fig. 4E). Interestingly, these BIM^hi^ cells largely overlapped with the BCL2^+^ population (Fig. 4F), suggesting a selective enrichment of a BCL2⁺BIM^hi^ IgM⁺ immature-like sub-state in *EμMyc Tet2*^−/−^ mice. These data point to a cell subset with increased BIM-dependent death pressure which is buffered by enhanced protection through BCL2.

Because expression changes and steady-state protein abundance do not necessarily predict survival dependence, we next evaluated spontaneous apoptosis in short-term culture. As reported previously, *EμMyc* IgM⁺ immature-like B cells rapidly undergo spontaneous apoptosis^53,55,56,71,72^, however, *Tet2* loss provided partial protection from death, most evident at 6 hours (Fig. 4G).

Given the prominent BCL2⁺BIM^hi^ fraction in *EμMyc Tet2*^−/−^ IgM⁺ immature-like B cells (Fig. 4F), we next assessed drug-induced changes in cytochrome c release, which reflect mitochondrial outer membrane permeabilization and thus commitment to intrinsic apoptosis. Splenic B cells from premalignant mice were treated for 1 hour with the BCL2 inhibitor ABT-199 (Venetoclax) or the MCL1 inhibitor S63845. ABT-199 induced greater cytochrome c release in *EμMyc Tet2*^−/−^ IgM⁺ immature-like B cells compared with *EμMyc* controls in this short-term *in vitro* priming assay (Fig. 4H), consistent with increased functional BCL2 dependence in this subset. In contrast, S63845 had little effect on cytochrome c release (Fig. 4I).

Altogether, these findings show the selective enrichment of an apoptosis-buffered IgM⁺ immature-like sub-state upon *Tet2* loss in the *EμMyc* background.

### *Tet2* loss enhances persistence and clonal skewing in premalignant IgM⁺ *EμMyc* B cells

Given the reduced apoptosis sensitivity in premalignant *EμMyc Tet2^−/−^* IgM⁺ immature-like B cells (Figure 4), we asked whether this compartment also contains cells with an improved capacity to persist and divide. Therefore, we tracked cell division of premalignant splenic B cells over 48 hours in culture via proliferation dye loss. Most IgM⁺ immature-like B cells were rapidly lost over 48 hours (Fig. 5A), consistent with the pronounced *in vitro* apoptosis sensitivity observed in Figure 4G. The few surviving cells in *EμMyc* cultures largely failed to divide, whereas a small fraction of *EμMyc Tet2^−/−^* IgM⁺ immature-like B cells persisted and underwent multiple divisions within 48 hours (Fig. 5B). To assess whether the dye dilution phenotype could be explained by increased proliferative capacity, we profiled DNA content *ex vivo* and performed a 2 hour *in vitro* 5-ethynyl-2’-deoxyuridine (EdU) pulse combined with phosphohistone H3⁺ (pH3) co-staining in premalignant B cells. DNA content analysis showed an increased fraction of IgM⁺ immature-like cells in the S and G_2_/M cell cycle phases (Fig. S5A, B). In line, EdU incorporation was moderately enhanced in *EμMyc Tet2^−/−^* IgM⁺ immature-like B cells, while the fraction of EdU⁺pH3⁺ cells, which reflects progression into mitosis among EdU-labeled cells, was comparable between genotypes (Fig. S5C, D). Of note, IgM⁻ progenitors showed no clear genotype-associated differences across these measures (Fig. S5A-D). Together, these data suggest improved persistence and modest redistribution across cell cycle phases, which could reflect altered cell cycle kinetics within the IgM⁺ immature-like compartment.

**Figure 5:**
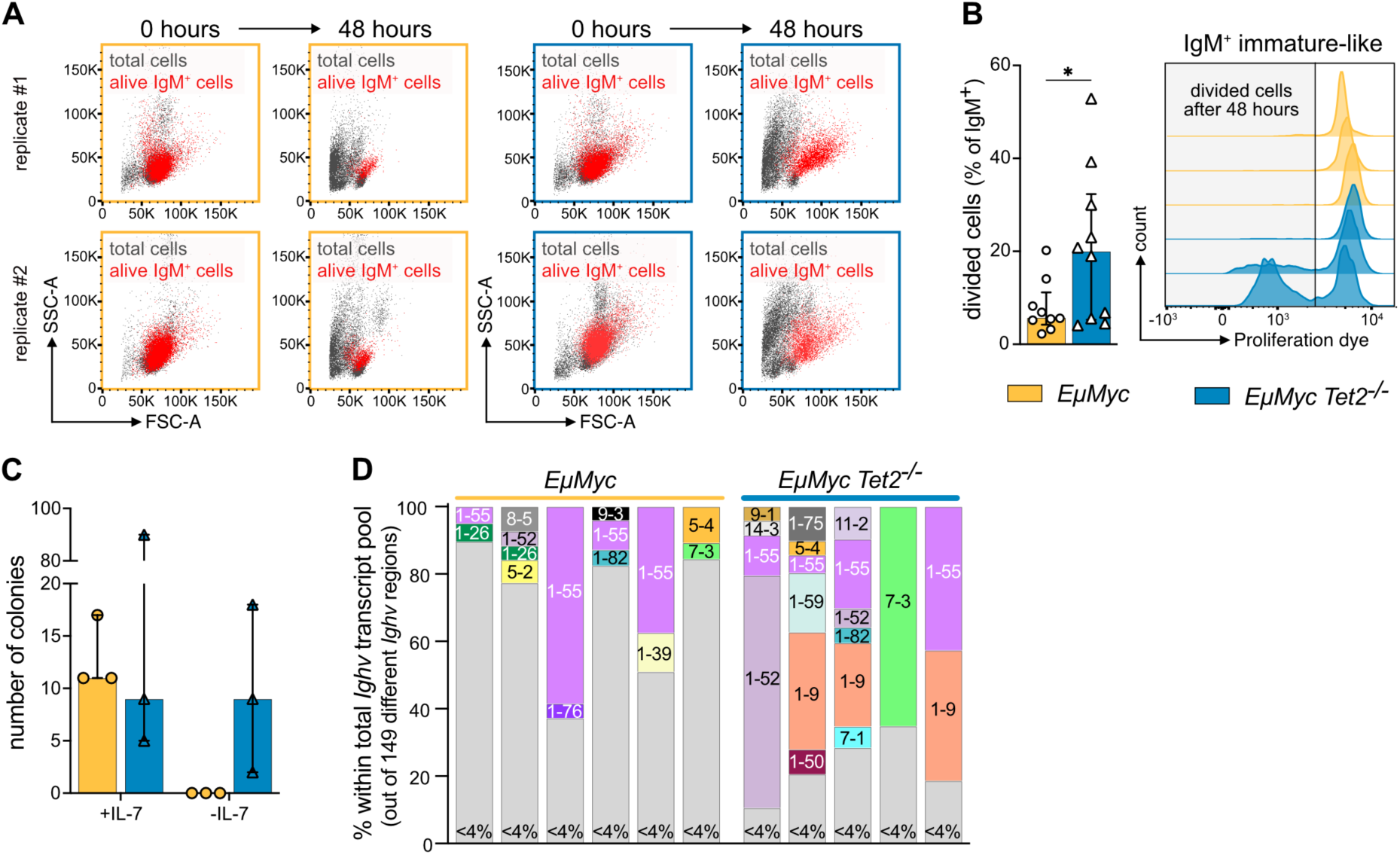
*Tet2* loss enhances persistence and clonal skewing in premalignant IgM⁺ *EμMyc* B cells. Analysis of proliferative capacity in splenic IgM^+^ immature-like B cells from *EµMyc* (n=9) and *EµMyc Tet2^−/−^* (n=10) mice. Total splenocytes were labeled *in vitro* with a proliferation dye, and flow cytometric analysis was performed at 0 and 48 hours. **(A)** Representative dot plots showing an overlay of total cells (grey) and live IgM^+^ immature-like B cells (red) at 0 and 48 hours. Two replicates of each genotype are depicted. **(B)** The fraction of dividing IgM^+^ immature-like B cells was quantified based on proliferation dye dilution. Representative flow cytometry histograms of proliferation dye intensity in IgM^+^ immature-like B cells from *EµMyc* (left) and *EµMyc Tet2^−/−^* (right) mice. **(C)** Splenocytes from premalignant *EµMyc* (n=3) and *EµMyc Tet2^−/−^* (n=3) were plated in methylcellulose with or without IL-7, and colonies were counted after 7 days. **(D)** Immunoglobulin heavy chain variable region (*Ighv*) gene usage in IgM^+^ immature-like B cells was determined by RNA-seq. Each bar represents an individual mouse, with *Ighv* genes comprising >4% of total *Ighv* transcripts displayed in distinct colors with corresponding gene names. Bar plots show median with interquartile range. Statistical significance was determined using (B) unpaired t-test. Normality was assessed using the Shapiro-Wilk test. *p<0.05

To test whether this phenotype associates with enhanced colony forming capacity, we plated splenocytes in methylcellulose with or without the cytokine IL-7, which is required for the survival of IgM⁻ progenitors but not IgM^+^ immature B cells^65,73^. Strikingly, *EμMyc Tet2^−/−^* cells showed an increased capacity to form colonies without IL-7 (Fig. 5C), indicating enhanced clonogenic potential. Consistent with a fitness advantage emerging in only a subset of premalignant IgM⁺ immature-like B cells, we next asked whether *Tet2* loss is accompanied by early clonal skewing within this compartment. Using immunoglobulin heavy chain variable region (*Ighv*) transcript abundance as a proxy for clonal representation, premalignant *EμMyc Tet2^−/−^* IgM⁺ immature-like B cells displayed a more restricted repertoire than *EμMyc* controls (Fig. 5D), whereas overall Variable Heavy chain (VH) gene usage patterns were broadly similar between genotypes (Fig. 5D). These data support the view of increased clonal skewing rather than a genotype-specific shift in VH preference. Consistent with repertoire compression, differential expression analysis in the same bulk RNA-seq dataset showed broad underrepresentation of immunoglobulin variable-region transcripts in *Tet2*-deficient IgM⁺ cells, supporting repertoire compression rather than uniform regulation of Ig gene expression (Fig. S5E).

Together, these data suggest that *Tet2* loss enables a small subset of premalignant IgM⁺ immature-like *EμMyc* B cells to persist and expand. This early fitness and clonal skewing likely contribute to the later increased representation of IgM⁺ lymphomas and the fully penetrant disease observed in *EμMyc Tet2^−/−^* mice.

## DISCUSSION

The gene encoding the DNA demethylase TET2 is recurrently affected by LOF mutations across a broad range of hematologic malignancies. However, how *Tet2* loss contributes to transformation remains incompletely understood. Here, we used the *EμMyc* mouse model to assess whether and how *Tet2* loss affects MYC-driven B cell lymphomagenesis. We show that TET2 deficiency increases lymphoma penetrance and shifts the tumor spectrum toward IgM⁺ disease, while once established, lymphomas appeared broadly similar when comparing *EμMyc* and *EμMyc Tet2^−/−^* mice. These findings suggest that *Tet2* loss acts mainly during premalignant stages of MYC-driven lymphomagenesis.

In mouse models, *Tet2* loss *per se* is only weakly tumorigenic and associates with long disease latency, suggesting a role as tumor facilitator rather than autonomous driver^12–15^. In B cells, direct evidence that *Tet2* loss cooperates with defined oncogenic drivers remains limited to a few studies, most notably TCL1A- and BCL6-driven malignancies^16,47^. This contrasts with mouse models of myeloid malignancies such as MDS, MPN-like disease, CMML, and AML, where the facilitator role is firmly established. Here, TET2 deficiency cooperates with lesions affecting signaling pathways and genome surveillance, including JAK2^V617F^, oncogenic KIT, FLT3-ITD, oncogenic RAS, and *Tp53* loss^14,38,42,46,60^. Notably, several of these alterations are themselves associated with MYC activation or MYC-associated transcriptional programs^74–76^. In the context of oncogenic KIT, TET2 deficiency has been linked to a hyperactive MYC signature downstream of PI3K signaling^60^. Consistent with this, *Myc*/*MYC* and *Tet2*/*TET2* transcripts are inversely related in MYC-driven mouse T-cell acute lymphoblastic leukemia (T-ALL) and in the human Burkitt cell line P493-6, where MYC also binds the *TET2* locus^77^. Beyond expression control, MYC can engage TET2 both at chromatin and through metabolism: in U2OS cells via SNIP1-dependent recruitment to chromatin, and in human and murine B lymphoma models through αKG/2-hydroxyglutarate-dependent modulation of TET activity and 5hmC states^78–80^. Against this background, our *EμMyc* model provides the first direct *in vivo* evidence that *Tet2* loss cooperates with oncogenic MYC in tumorigenesis. This is particularly relevant in B cells, where MYC alterations are prominent drivers of lymphoma development^81,82^.

One of the most striking phenotypes in *EμMyc Tet2^−/−^*mice is the shift toward IgM⁺ tumors. In premalignant mice, this is mirrored by developmental skewing at the transition from immature to mature B cells, making IgM⁺ immature-like B cells the key population. This compartment was less homogeneous than expected, and our data suggest that *Tet2* loss enriches distinct substates within it. The strongly reduced CD21⁺ and CD23⁺ cell frequencies in premalignant *EμMyc Tet2^−/−^* mice place the enriched substates toward the less mature end of the IgM⁺ immature-like compartment. Consistent with this, we do not detect appreciable *Aicda* or *Bcl6* expression, in line with prior work indicating that these cells retain an immature state^58,59^.

Excluding immunoglobulin genes, which likely reflect clonal skewing, GO-term enrichment for lymphocyte differentiation, negative regulation of cell development, and regulation of hemopoiesis was evident in *EμMyc Tet2^−/−^* IgM⁺ immature-like B cells. Together with reduced inferred HNF1A and TCF3/E2A activity by decoupleR, this is more consistent with impaired progression toward mature B cells rather than with a strong developmental block^83,84^. Related perturbations of differentiation have likewise been described upon *Tet2* loss in other hematopoietic contexts, including HSPCs^13–15^ and in germinal center (GC) B cells^24^.

Apart from the CD21/CD23 phenotype, increased CD9⁺, CD80⁺, and CD86⁺ cell fractions indicate substates with enhanced interaction and activation potential. This is supported by GO-term enrichment for negative regulation of cell activation, regulation of T cell activation, and immune response-regulating signaling, together with increased inferred NR4A1, KLF4, JUNB, and RUNX1 activity by decoupleR. CD86 and CD9 are best supported by prior literature. In mice, B cell-specific TET loss caused *Cd86* de-repression via reduced HDAC1/2 recruitment and altered chromatin at the *Cd86* locus^22^. In murine STAT5-driven leukemic stem-cell models, CD9 marks a high-fitness, self-renewing subpopulation linked to JAK/STAT-associated persistence under oncogenic stress^85^. In our data, enrichment of survival-promoting JAK/STAT, TNF/NF-κB, and related signaling programs likely reflects a broader logic also seen in myeloid models, in which *Tet2* loss repeatedly shifts signaling responsiveness, stress handling, and competitive fitness toward persistence under oncogenic pressure^14,38,42,46,60^. This suggests that, under oncogenic MYC, premalignant B cells use *Tet2* loss in a manner that parallels what has been observed in oncogene-driven myeloid cells.

Under strong MYC-imposed apoptotic pressure, such signaling states are well positioned to favor survival of cells that can better buffer mitochondrial death signaling. Notably, *EμMyc Tet2^−/−^* mice selectively enriched a BCL2⁺BIM^hi^ IgM⁺ immature-like subpopulation that represents only a minor fraction in *EμMyc* controls. The *EμMyc* model is characterized by MYC-induced BIM and reduced BCL2 in IgM⁺ B cells, and by preferential expansion of these cells upon BIM loss^56^. Such a BCL2⁺BIM^hi^ population fits a scenario in which BIM-associated apoptotic priming is buffered by BCL2, thereby promoting cell survival^86–92^. Accordingly, *EμMyc Tet2^−/−^* IgM⁺ immature-like cells showed a modest but reproducible survival advantage during spontaneous apoptosis *in vitro*, reflecting increased fitness under conditions of growth factor withdrawal and loss of microenvironmental interactions. The BCL2-inhibitor ABT-199, but not the MCL1-inhibitor S63845, increased cytochrome c release in these cells, consistent with enrichment of a functionally BCL2-dependent BCL2⁺BIM^hi^ substate.

Beyond our system, evidence linking *Tet2* loss to BCL2-family dependencies and cell survival remains limited. In B cells, the clearest survival-related evidence comes from a murine CLL model, in which *Tet2* loss was associated with stronger BCR signaling dependency^16^. In *EμMyc Tet2^−/−^* IgM⁺ immature-like B cells, however, the phospho-readouts do not support enhanced tonic or proximal BCR signaling. Although pAKT was increased, this readout is not specific for BCR activity and is equally compatible with broader survival-associated signaling. In HSPC and myeloid contexts, *Tet2* loss is most consistently linked to enhanced self-renewal, competitive fitness, and resistance to inflammatory stress rather than to broad proliferative activation^5,42,93^. Along this line, premalignant *EμMyc Tet2^−/−^* IgM⁺ immature-like B cells, aberrantly driven into cycle by MYC, showed only a modest shift toward S and G_2_/M and no consistent transcriptomic evidence of enhanced proliferation. Yet after 48 hours in culture, the few surviving *EμMyc Tet2^−/−^*IgM⁺ immature-like B cells had proliferated, unlike their *EμMyc* counterparts. Further, *EμMyc Tet2^−/−^* IgM⁺ immature-like B cells showed increased colony-forming capacity in methylcellulose and evidence of repertoire compression and clonal skewing based on *Ighv* transcript abundance. Overall, our data suggest that *Tet2* loss promotes persistence and selective outgrowth rather than broad proliferative activation, extending to premalignant B cells a principle previously established in TET2-deficient HSPC and myeloid settings.

Several aspects of the present study also define the limits within which this model should be interpreted. Here, the *EμMyc* system is used as a mechanistic model of MYC-driven lymphomagenesis rather than a preclinical surrogate for a single human lymphoma entity. In addition, our data suggest that heterogeneity within the premalignant IgM⁺ compartment has been underappreciated in *EμMyc* mice. While our bulk analyses do not fully resolve this heterogeneity, they still provide consistent transcriptomic, phenotypic, and functional evidence that *Tet2* loss has biologically meaningful effects in this compartment. Finally, because *Tet2* loss was analyzed in a germline setting, B cell-intrinsic and non-cell-autonomous effects cannot be fully separated, although this may also reflect biologically relevant aspects of early hematopoietic *Tet2* loss, including clonal hematopoiesis. More broadly, it will be important to determine whether similar premalignant survival states also arise in more human-relevant hematopoietic disease settings. Even within these constraints, the main conclusions remain unchanged.

Using the *EμMyc* model, we provide a mechanistic framework for how *Tet2* loss facilitates MYC-driven B cell lymphomagenesis through apoptosis buffering and selective clonal outgrowth. Together, these findings support a model in which TET2 deficiency promotes lymphoma development by enhancing survival under MYC-imposed stress and thereby increasing the likelihood of premalignant clonal persistence, selection and outgrowth.

## MATERIALS & METHODS

### Animal models

All animals were backcrossed and maintained on a C57BL/6N background for at least 10 generations and bred at the central laboratory animal facility of the Medical University of Innsbruck. Animal experiments were approved by the Austrian Federal Ministry of Education, Science and Research (BMWF: 66.011/0008-V/3b/2019) and conducted under standard housing conditions consisting of a 12-hour (h) light/dark cycle, relative humidity of 55-65% and temperature 22 ± 2°C.

*EµMyc* (B6.Tg(IghMyc)22Bri) and *Tet2^−/−^* (B6(Cg)-*Tet2^tm^*^1^*^.2Rao^*) mouse lines were generated and genotyped as previously described^13,55^. Both male and female mice were used indiscriminately and were monitored until either 46 - 55 days of age, 280 - 320 days of age, or until meeting predetermined euthanasia criteria.

### Preparation of single-cell suspensions

All centrifugation steps were performed at 1500 rpm for 5 minutes (min) at 4°C (VWR Centrifuge Megastar 1.6R).

Bone marrow cells were isolated by flushing femurs and tibiae with FACS-B buffer (PBS with 2% FBS (Gibco, 10270106) and 10 µg/ml Gentamicin (Gibco, 15750037)) using a syringe and a 23G needle. The cell suspension was centrifuged and filtered through a 50 µm filter (BD Bioscience, 340632).

Spleens were dissociated by gently pressing the tissue through 70 µm cell strainers (Corning, 352350). The resulting cell suspension was centrifuged and resuspended in 1 ml of ice-cold red blood cell lysis buffer (155 mM NH4Cl, 10 mM KHCO3, 0.1 mM EDTA; pH 7.5) and incubated on ice for 3 min. Lysis was stopped by adding 5 ml of FACS-B, centrifuged and filtered through a 50 µm filter.

Cell numbers in suspension were determined using hemocytometer and trypan blue exclusion.

### Cell surface staining for flow cytometry

Splenocytes and bone marrow cells were stained for flow cytometric analysis, with a minimum acquisition threshold of 3*10^5^ cells per sample. All centrifugation steps were performed at 2000 rpm for 2 min at 4°C (VWR Centrifuge Megastar 1.6R), and all incubations were conducted at 4°C in 96-U-well plates. Cells were incubated for 10 min with 20 µl of αCD16/32 Fc-Block (1:200 in FACS-B; Biolegend, 101310), followed by incubation for 15 min with 30 µl of a mixture of the following fluorochrome-conjugated anti-mouse antibodies diluted in FACS-B supplemented with Brilliant Stain Buffer (1:4; BD Biosciences, 566349): αCD9-FITC (1:400, Biolegend, 124808), αCD40-FITC (1:200, eBioscience, 11-0402-81), αCD8-PE (1:300, Biolegend, 100708), αNK1.1-PeCy7 (1:100, Biolegend, 108713), αCD86-PeCy7 (1:100, Biolegend, 105013), αIgM-PeCy7 (1:200, Biolegend, 406514), αIgD-PerCP/Cy5.5 (1:200, Biolegend, 405710), αCD38-APC (1:200, Biolegend, 102712), αIgM-APC (1:1000, Jackson ImmunoResearch, 115-607-020), αCD4-A700 (1:400, Biolegend, 116022), αCD21-A700 (1:200, Biolegend, 123431), αCD23-APC/Cy7 (1:200, Biolegend 101629), αGr1-BV421 (1:200, Biolegend, 108433), αCD80-BV421 (1:100, Biolegend, 104725), αCD5-BV421 (1:200, Biolegend, 100617), αTer119-BV510 (1:100, Biolegend, 116237), αCD44-BV605 (1:200, Biolegend, 103047), αTCRβ-BV605 (1:200, Biolegend, 109241), αCD11b-BV650 (1:1000, Biolegend, 101259), αCD19-BV711 (1:400, Biolegend, 115555) and αB220-BV785 (1:400, Biolegend, 103246). Finally, cells were washed with FACS-B and immediately acquired on an LSR II-Fortessa flow cytometer (BD Bioscience). Data were analyzed using FlowJo software (version 10.10.0).

### Intracellular staining for flow cytometry

All centrifugation steps were performed at 2400 rpm for 2 min at 4°C (VWR Centrifuge Megastar 1.6R), and all incubation steps were carried out at 4°C in 96-U-well plates. Following cell surface staining (see section *Cell surface staining*), cells were fixed and permeabilized by incubation with 100 µl FixPerm buffer (containing 4% methanol-free formaldehyde (Thermo Fisher, 28908) and 0.1% Saponin (Sigma-Aldrich, 47036)) for 20 min and then washed three times with 150 µl PermWash buffer (PBS with 1% BSA, 0.1% Saponin, 0.0025% Natrium Azid).

For DNA content analysis, in combination with mitotic and DNA damage markers, cells were incubated with 30 µl αCD16/32 Fc-Block (1:100 in PermWash Buffer) for 15 min, followed by incubation with 30 µl of a mixture of the following fluorochrome-conjugated anti-mouse antibodies for at least 30 min: αpH3-PE (1:100, Biolegend, 650807) or cleaved Caspase-3-PE (1:100, BD Bioscience, 570183) and αγH2AX-PerCPCy5.5 (1:100, eBioscience, 46-9865-42). Cells were washed with 100 µl PermWash Buffer and incubated in 100 µl of PBS containing 250 µg/ml RNase A (Sigma-Aldrich, R5500) at 37° for 20 min, after which 50 µl of TO-PRO-3 Iodide (1:333 in PBS, Thermo Fisher, T3605) was added and samples were immediately acquired.

For analysis of BCR signaling components, cells were incubated with 30 µl αCD16/32 Fc-Block (1:100 in PermWash Buffer) for 15 min, followed by incubation with 30 µl of following primary antibodies for at least 30 min: B cell Signaling Antibody Sampler (1:100, Cell Signaling, 9768), αpAKT (1:100, Cell Signaling, 4060S), αAKT (1:100, Cell Signaling, 4691S), αSYK (1:100, Santa Cruz, sc1077). After a washing step with 100 µl PermWash Buffer, cells were incubated with a goat anti-rabbit IgG (H+L) Alexa Fluor^TM^ 647-conjugated secondary antibody (1:1000, Invitrogen, A21245) for 15 min at 4°C. Finally, cells were washed with 100 µl FACS-B and acquired immediately.

To assess BCL2 family members, cells were incubated with 30 µl αCD16/32 Fc-Block (1:20 in PermWash Buffer) for 15 min, followed by incubation with 30 µl of following antibodies: αBIM (1:100, Abcam, ab32158), αBCL2-PE (1:100, Biolegend, 633507), αMCL1 (1:100, Cell Signaling, 5453T) or αBCL-XL (1:100, Cell Signaling, 2764S) for at least 30 min. After a washing step with 100 µl PermWash Buffer, a goat anti-rabbit IgG (H+L) Alexa Fluor^TM^ 647-conjugated antibody (1:1000, Invitrogen, A21245) was applied for 15 min. A final washing step with 100 µl PermWash Buffer was performed.

Samples were immediately acquired on an LSR II-Fortessa flow cytometer (BD Bioscience), and data were analyzed using FlowJo software (version 10.10.0).

### B cell viability assay

B cells were enriched from splenic cell suspensions using MagniSort^TM^ Streptavidin Negative Selection Beads (ThermoFisher, MSNB-6002-74), according to the manufactureŕs instruction. For depletion of non-B cells, 300 µl of a biotinylated antibody mix (diluted 1:100 in FACS-B) containing αCD4 (Biolegend, 100404), αCD8 (Biolegend, 100704), αNK1.1 (Biolegend, 108704), αCD11b (Milteny Biotec, 130-113-242), αGr1 (Biolegend, 108404) and αTer119 (Biolegend, 116204) was used.

B cells (5*10^5^) were then cultured in 50 µl FACS-B (untreated) in 96-U-well plates for 0 h, 2 h, 6 h and 10 h at 37°C. To avoid loss of dead cells, 25 µl of the following antibody mix (diluted in FACS-B) was added directly to each well: αCD16/32 Fc-Block (1:67; Biolegend, 101310), αCD19-PE (1:67; eBioscience, 12-0191-83), αB220-PeCy7 (1:67; Biolegend, 103222), αIgD-PerCP/Cy5.5 (1:200, Biolegend, 405710), αIgM-APC (1:1000, Jackson ImmunoResearch, 115-607-020) and DAPI (1:16600; Sigma-Aldrich, D9542). Samples were acquired on a LSR II-Fortessa flow cytometer (BD Bioscience), and data were analyzed using FlowJo software (version 10.10.0).

### Drug-induced cytochrome c release

Apoptotic sensitivity was assessed based on a previously published protocol^88,89,94^. Briefly, 30 µM ABT-199/Venetoclax, (MedChemExpress, HY-15531) 1 µM S63845 (MedChemExpress, HY-100741), 20 µM Alamethicin (MedChemExpress, HY-N6708) and 1% dimethyl sulfoxide (DMSO; Merck, D5879) were diluted to 2X the desired final concentration in 25 µl of 0.002% digitonin (Merck, D141) prepared in MEB buffer (Mannitol Experimental Buffer: 10 mM HEPES (Merck, H0887) pH 7.5, 150 mM mannitol (Merck, M9647), 150 mM KCl, 1 mM EGTA (Merck, E3889), 1 mM EDTA (Merck, ED4S), 0.1% BSA (Sigma-Aldrich, 12659), 5 mM succinate (Merck, S3674)). These compound plates were pre-arrayed in 96-U-well plates and stored at −80°C until use.

Per treatment, 2*10^5^ splenocytes were stained (see section *Cell surface staining*) in low-binding tubes (Sarstedt, 72706600) with 100 µl of the following antibodies diluted in FACS-B supplemented with Brilliant Stain Buffer (1:4; BD Biosciences, 566349): αCD16/32 (1:200, Biolegend, 101310), αIgM-PeCy7 (1:200, Biolegend, 406514), αIgD-PerCP/Cy5.5 (1:200, Biolegend, 405710), αCD19-BV711 (1:400, Biolegend, 115555) and αB220-BV785 (1:400, Biolegend, 103246). Cells were subsequently washed with FACS-B and further incubated with 300 µl Zombie Green fixable viability dye (1:500 in FACS-B; Biolegend 423111) for 10 min at 4°C. Per well 130 µl FACS-B was added and centrifuged at 2000 rpm for 2 min at 4°C (VWR Centrifuge Megastar 1.6R). For each 96-U-well, 2*10^5^ stained cells (in 25 µl MEB buffer) were added to 25 µl compound solution (pre-arrayed compound plates were thawed 1 hour beforehand), and incubated for 1 h at RT in the dark. Subsequently, 16.5 µl of 4% methanol-free formaldehyde (Thermo Fisher, 28908) was added per well and incubated for 10 min at RT, followed by the addition of 16.5 µl per well N2 buffer (1.7 M Tris base (Carl Roth, 5429.4), 1.25 M Glycin (Fisher Scientific, 10773644); pH 9.1) and incubation for 5 min at RT. Finally, 10 µl per well anti-cytochrome c antibody (1:400, Biolegend, 612310) diluted in 10x CytoStain Buffer (PBS with 2% Tween 20 (Carl Roth, 9127.1) and 10% BSA) was added and incubated for 12 h at 4°C in the dark before flow cytometric analysis on an LSR II-Fortessa flow cytometer (BD Bioscience). Data were analyzed using FlowJo Software (version 10.10.0).

### Colony formation assay (MethoCult)

Colony-forming unit assays using MethoCult were performed according to the manufactureŕs instructions. Briefly, splenocytes (1*10^5^) were resuspended in 100 µl IMDM (Gibco, 21056023), supplemented with 100 units/ml penicillin and 100 µg/ml streptomycin (Sigma-Aldrich, P0781), and plated in 35-mm culture dishes (TC Dish35, Suspension, Sarstedt, 83.3900.500) in 1 ml methylcellulose without IL-7 (MethoCult M3231, STEMCELL Technologies) or with IL-7 (MethoCult M3630, STEMCELL Technologies). Dishes were incubated at 37°C, and colonies were counted after 7 days.

### *In vitro* 5-ethynyl-2-deoxyuridine (EdU) assay

Splenocytes (4*10^5^) were incubated with 10 µM EdU using the Click-iT^TM^ Plus EdU Flow Cytometry Assay Kit (Invitrogen, C10632) for 2 h according to the manufactureŕs instructions. Following EdU labeling, cells were harvested and washed twice with 200 µl FACS-B.

Cell surface staining was subsequently performed (see section *Cell surface staining*) using the following fluorochrome-conjugated anti-mouse antibodies diluted in FACS-B supplemented with Brilliant Stain Buffer (1:4; BD Biosciences, 566349): αCD16/32 (1:200, Biolegend, 101310), αIgM-FITC (1:200, Biolegend, 406505), αIgD-PerCP/Cy5.5 (1:200, Biolegend, 405710), αCD19-BV711 (1:400, Biolegend, 115555), and αB220-BV785 (1:400, Biolegend, 103246). Following surface staining, cells were fixed, permeabilized, and stained as described in section *Intracellular staining for flow cytometry* with αpH3-PE (1:200, Biolegend, 650807). EdU detection was then performed according to the manufactureŕs instructions. Samples were acquired on an LSR II-Fortessa flow cytometer (BD Bioscience) and data were analyzed using FlowJo software (version 10.10.0).

### Proliferation assay

Splenocytes (2*10^6^) were labeled with 10 µM Cell Proliferation Dye eFlour^TM^ 450 (Thermo Fisher, 65-0842-90) according to the manufactureŕs instructions. Briefly, cells were harvested and washed twice with 200 µl pre-warmed PBS (centrifugation: 2000 rpm for 2 min at 24°C; Eppendorf centrifuge 5424R) to remove serum. A 20 µM dye solution was prepared in pre-warmed PBS and mixed 1:1 with the cell suspension while vortexing. The mixture was incubated for 10 min at 37°C in the dark. Labeling was stopped by adding 4 volumes of cold complete B cell medium (DMEM medium (Sigma-Aldrich, D6429) supplemented with 10% FBS (Gibco, 10270106), 2 mM L-glutamine (Sigma-Aldrich, G7513), 50 µM ß-mercaptoethanol (Sigma-Aldrich, M3148), 10 mM HEPES (Sigma-Aldrich, H0887), 1 mM sodium pyruvate (Gibco, 11360039), 1X nonessential amino acids (Gibco, 11140035), 100 units/ml penicillin and 100 µg/ml streptomycin (Sigma-Aldrich, P0781)), followed by incubation on ice for 5 min. After three washing steps with 1 ml complete B cell medium (centrifugation: 2000 rpm for 2 min at 24°C; Eppendorf centrifuge 5424R), labeled cells were cultured in 500 µl B cell medium per well in 24-well plates. Following 48 h of culture, cells were gently resuspended, transferred to a 96-U-well plate, and stained (see section *Cell surface staining*) with 30 µl of the following fluorochrome-conjugated anti-mouse antibodies diluted in FACS-B: αIgD-PerCP/Cy5.5 (1:200, Biolegend, 405710), αIgM-APC (1:1000, Jackson ImmunoResearch, 115-607-020), αCD19-BV711 (1:400, Biolegend, 115555), and αB220-BV785 (1:400, Biolegend, 103246). After cell surface staining, cells were incubated in 100 µl PBS containing fixable viability dye (1:2000, eBioscience 65-0866-14) for 10 min at 4°C and washed with 130 µl FACS-B. Samples were resuspended in FACS-B buffer and acquired on an LSR II-Fortessa flow cytometer (BD Bioscience). Data were analyzed using FlowJo Software (version 10.10.0).

### Cell sorting

Splenocytes were incubated for 15 min at 4°C in 300 µl of the following fluorochrome-conjugated anti-mouse antibodies: αB220-FITC (1:200, eBioscience, 553088), αCD25-PE (1:200, Biolegend, 101904), αIgD-PerCP/Cy5.5 (1:200, Biolegend, 405710), αIgM-APC (1:1000, Jackson ImmunoResearch, 115-607-020), αCD117(c-kit)-BV421 (1:200, Biolegend, 105828), and αCD19-BV605 (1:200, Biolegend, 115540). Subsequently, 1 ml of FACS-B was added, cells were centrifuged (1500 rpm for 5 min at 4°C; VWR Centrifuge Megastar 1.6R), resuspended in 0.5-3 ml FACS-B, and filtered through a 50 µm cell strainer (BD Bioscience, 340632).

Cell sorting was performed at 4°C on a FACS Aria III (70 µm nozzle; BD Bioscience). Doublets were excluded based on FSC-H/FSC-W and SSC-H/SSC-W gating. Sorted cell subsets were defined as follows: large pre-B cells (B220^+^CD19^+^IgM^−^IgD^−^cKit^−^CD25^+^FSC-A^hi^), IgM^−^ progenitor B cells (B220^+^CD19^+^IgD^−^IgM^−^), IgM^+^ (immature) B cells (B220^+^CD19^+^IgD^−^IgM^+^), and tumor cells (B220^+^CD19^+^IgD^−^IgM^−^ or B220^+^CD19^+^IgD^−^IgM^+^). Cells were collected into low-binding tubes (Sarstedt, 72706600) containing 300 µl FACS-B. Sorted cells were centrifuged (2200 rpm for 2.5 min at 4°C; Eppendorf centrifuge 5424R), resuspended in 500 µl PBS, and centrifuged again. Resulting cell pellets were either snap-frozen directly or resuspended in 100 µl of DNA/RNA Shield (Zymo Research, R1200-25) and stored at −80°C.

### RNA extraction, sequencing, and data analysis

Total RNA was extracted from frozen samples (pellets or in DNA/RNA Shield) using the Quick-RNA Micro Prep Kit (Zymo Research, R1050) with DNase digestion according to the manufactureŕs instructions. Library preparation and bulk RNA sequencing were performed using poly(A) selection, strand-specific library preparation and sequencing on an Illumina® NovaSeq^TM^ platform with 2×150 bp paired-end reads, targeting 20 million reads per sample. Data quality was guaranteed to be ≥85% bases with Q30 or higher.

Raw FASTQ reads were processed using the nf-core RNA-seq pipeline^95^ with the STAR-salmon^96^ option and aligned to the mouse reference genome GRCm39 (version M30). Genes with names. Starting with “Gm” or “Rp” or ending. With “rik” were not. Considered for the downstream analysis. In addition, genes were required to have at least 5 raw counts in at least “M” samples, where “M” was the size of the smallest group considered in the comparison. DEG analysis was performed using the DESeq2^97^ (version 1.44.0) package in RStudio (version 2024.12.0+467). Unless otherwise stated, genes were considered significantly differentially expressed (DEG) if they met the criteria of p-adjusted<0.05 and absolute log_2_(fold change)>1. For visualization of the B-lineage regulatory/maturation (Fig. 3C) and BCL2 family gene panel (Fig. 4B), volcano plots were generated using relaxed significance thresholds (p-adjusted<0.05 and absolute log_2_(fold change)>0.5) to highlight biologically relevant genes with modest expression changes. Volcano plots displayed all detected genes, except for the tumor cell comparison (Fig. 1F), in which immunoglobulin (*Ig*) genes were excluded to improve visualization clarity.

The B-lineage regulatory/maturation gene panel comprised 103 genes, including 13 upregulated genes (*Blk, Cd5, Cd9, Cd38, Cd44, Cd80, Cd83, Lef1, Myb, Pim2, Rbpj, Syk, Tcf4*), 14 downregulated genes (*Aicda, Bach2, Bcl6, Cd40, Cd93, Cebpa, Cr2, Ebf1, Fcer2a, Irf8, Lmo2, Slamf6, Slamf1, Zap70*) and 76 genes not significantly deregulated (*Actb, Bank1, Bcl11a, Blnk, Btk, Ccnd3, Ccr7, Cd14, Cd19, Cd22, Cd24a, Cd27, Cd28, Cd48, Cd53, Cd69, Cd72, Cd74, Cd79a, Cd79b, Cd81, Cd86, Cd200, Cd300a, Cxcr4, E2f1, Ets1, Fcer1g, Foxo1, Foxp1, Foxp4, Ikzf1, Ikzf2, Ikzf3, Il7r, Irf4, Itga4, Itgam, Itgb2, Lcp2, Ly9, Lyn, Mafb, Mcl1, Ms4a1, Myc, Notch2, Pax5, Paxip1, Plcg2, Ppp3cb, Prdm1, Ptpn6, Rag1, Rag2, Rela, Runx1, Smad3, Spi1, Spib, Stat5a, Stat6, Tcf3, Tnfrsf13b, Tnfrsf13c, Tnfrsf17, Vav1, Vav2, Vpreb1, Xbp1, Zbtb7a, Zbtb7b, Zfp386, Zfp423, Zfp521, Zyx*).

The BCL2 family gene panel consisted of 17 genes, including 6 upregulated genes (*Bbc3, Bcl2, Bcl2l11, Bcl2a1b, Bcl2a1d, Blk*), 1 downregulated gene (*Bmf*), and 10 genes not significantly deregulated (*Bcl2l1, Bcl2l2, Mcl1, Bad, Bid, Bak1, Bax, Pmaip1, Hrk, Bik*).

GO-term enrichment analysis for biological processes was performed on DEGs using the clusterProfiler package^98^ (version 4.12.6) in RStudio (version 2024.12.0+467). Enrichment analysis for MSigDB Hallmark genes was conducted using Enrichr^99–101^. Transcription factor activity was inferred using the decoupleR package^63^ (version 2.9.7) based on CollecTRI regulons^102^ in RStudio (version 2024.12.0+467) with a minimum threshold of 45 target genes per transcription factor and a significance cutoff of p-value<0.05.

Immunoglobulin heavy chain variable (*Ighv*) transcript usage is based on transcript per million (tpm) reads. The relative proportion of each *Ighv* gene was calculated by normalizing to the total *Ighv* transcript abundance (set to 100%). Individual *Ighv* transcripts exhibiting >4% were highlighted by individual color and gene name in bar plots, with each bar representing a single mouse.

The aneuploidy score shown in Figure S1G was calculated as follows: chromosome-wide copy-number profiles were inferred from bulk RNA-seq (premalignant and malignant) using the gene-dosage signal in differential expression with the R package DESeq2^97^ (version 1.50.2). For each sample/contrast, gene-level log_2_(fold change) estimates and standard errors (lfcSE) were mapped to genomic coordinates using GRCm38 Ensembl mouse gene annotations (chromosomes 1–19 and X). Genes with high uncertainty were removed (lfcSE ≥ 5). To place samples on a comparable baseline, gene-level log_2_(fold changes) were centered to an inferred euploid reference by computing a weighted mean log_2_(fold change) per chromosome with weights = 1/lfcSE, estimating the mode of the chromosome-mean distribution by kernel density, and subtracting this mode from all gene-level values within the sample. Smooth chromosome profiles were then obtained by first computing the running mean of weighted fold changes (window size 100, including partial sizes at both ends) and then fitting weighted generalized additive models separately for each chromosome and sample using the R package mgcv^103^ (version 1.9-3). Model predictions were evaluated on an evenly spaced genomic grid (∼10 Mb spacing) to generate genomic tiles, and predicted log2 fold-changes were converted to inferred absolute copy number compared to the diploid baseline.

### AI-based literature discovery

The Elicit web application (Pro subscription; https://elicit.com; accessed 2026) was used to support literature discovery by generating topic-focused research reports. All cited sources used for the manuscript were manually screened and verified by the authors. All prompts relevant to this manuscript are reported below in full.

06.02.2026: “Which mouse hematologic malignancy models show that Tet2 loss-of-function facilitates oncogene- or tumor-suppressor–driven disease (accelerates latency and/or increases penetrance) in myeloid and lymphoid lineages (B cell and T cell)? Primary outcomes (vs driver-only controls): incidence/penetrance (%) and latency/median survival (time-to-disease). Secondary outcomes (if reported): disease severity/progression (e.g., transformation, grade, leukocytosis/splenomegaly), and transcriptomic changes from RNA-seq (e.g., gene set enrichment / pathway signatures / MYC target modules, not just “DE genes”).” 06.02.2026: “In mouse or human hematopoietic cells and hematologic malignancy models, what direct functional evidence shows that TET2 loss-of-function impairs tumor-suppressor mechanisms in (1) DNA damage/repair, (2) apoptosis, (3) proliferation/cell-cycle control, (4) p53 signaling, (5) JAK–STAT signaling, (6) IL-6 signaling, (7) IFN signaling, (8) TNF signaling, or (9) NF-κB signaling? Include myeloid, B-cell, and T-cell contexts. Prioritize perturbation-based studies (TET2 loss ± rescue and/or pathway perturbation) with functional readouts. Primary outcomes (functional assays): DDR/repair/genomic instability: γH2AX/53BP1 foci, comet, micronuclei, karyotype/WGS/instability metrics Apoptosis/BCL2-family: BH3 profiling, Annexin V, caspases, mitochondrial priming/cytochrome c assays Proliferation/cell-cycle: EdU/BrdU, Ki-67, cell-cycle profiling p53 signaling: p53 activation/targets, checkpoint responses, genetic or pharmacologic modulation JAK–STAT / IL-6: pSTAT readouts, IL-6 dependence, inhibitor/rescue experiments IFN: ISG induction, IFN stimulation/blockade with functional consequences TNF / NF-κB: TNF stimulation/blockade, NF-κB activation/readouts with functional consequences”

04.02.2026: “In the EµMyc mouse model, what peer-reviewed evidence shows that proliferating/tumor-associated IgM⁺ IgD⁻ B cells are immature B cells (e.g., transitional/immature stages) rather than mature activated or memory B cells? Please extract the immunophenotyping panels and gating (e.g., B220, CD93/AA4.1, CD23, CD21, CD24, CD38, GL7, Fas, CD73), tissue location (BM vs spleen/LN), functional assays (transplantation/outgrowth/clonality), and molecular evidence (IgV SHM, AID signatures, BCR rearrangements) supporting the authors’ conclusion.”

04.02.2026: “In hematologic cells and tumors (mouse models and human systems), what evidence shows functional or genetic synergy between TET2 loss (KO, LOF mutation, knockdown) and BCL2-family anti-apoptotic proteins (e.g., BCL2, BCL-XL/BCL2L1, MCL1, BCL-W, A1/BCL2A1)? Focus on B-cell lymphomas, with a special emphasis on the EµMyc mouse model. Please extract the model/system, perturbations, readouts (tumor latency/burden, survival, apoptosis/BH3 profiling, drug sensitivity to BH3 mimetics), and whether the interaction is additive vs synergistic.”

09.03.2026: “In human and mouse hematologic malignancies, especially MYC-driven B-cell malignancies, which primary studies describe viable BIM-high, apoptosis-primed states that persist through buffering by BCL2 family proteins? Focus on premalignant or developmentally defined cell populations and on studies using functional mitochondrial apoptosis assays. If available, include evidence from TET2-mutant or TET2-deficient settings.”

09.03.2026: “In human and mouse hematologic malignancies, especially B-cell malignancies, which primary studies show that high BCL2 expression or functional BCL2 dependence is associated with increased sensitivity to selective BCL2 inhibition (for example venetoclax/ABT-199), particularly in apoptosis-primed cells with elevated BIM or other BH3-only proteins, and what evidence exists for this relationship in TET2-mutant or TET2-deficient settings?”

### Quantification and statistical analysis

Data are presented as median values with interquartile range. Normality of distributions was assessed using the Shapiro-Wilk test (α=0.05). For normally distributed data, statistical significance was evaluated using one-way ANOVA, two-way ANOVA, or an unpaired t-test, as appropriate. For non-normally distributed data, the Mann-Whitney test was applied. The Holm-Šidák method was used to correct for multiple comparisons where applicable. Data normalization procedures, when applied, are described in the corresponding Material and Methods subsections or in the figure legends. Significance thresholds were defined as *p<0.05, **p<0.005, ***p<0.0005, and ****p<0.0001. The specific statistical test and the number of biological replicates (n) for each analysis are indicated in the corresponding figure legends. Graphs and statistical analysis were performed using GraphPad Prism (version 10.6.0) and Microsoft Excel (version 16.78), and figures were created using Affinity Designer (version 1.10.8) and RStudio (version 2024.12.0+467).

## ACKNOWLEDGEMENTS

We thank Emmanuel Derudder, William Olson, Sebastian Herzog and Felix Eichin for insightful discussions. We are grateful to B. Unterberger, C. Soratroi and I. Gaggl for expert technical assistance and M. Saurwein for animal care. The AI-assisted web app tool Elicit was used for literature discovery, the details are provided in the Methods section. This research was funded in whole by the Austrian Science Fund (FWF) (Grant DOI 10.55776/FG25, “BCL2 Network Adaptations in B Cell Transformation”, to V.L., A.V., J.S.R. and F.F.). M.E. received funding from Deutsche Forschungsgemeinschaft (DFG) (Grant DOI TRR 353/1 - 471011418 – “Regulation of cell death decisions). F.F. was supported by the Austrian Science Fund (FWF) (Grant DOI 10.55776/PAT5895324). For open access purposes, the author has applied a CC BY public copyright license to any author accepted manuscript version arising from this submission. The computational results presented here have been achieved in part using the LEO HPC infrastructure of the University of Innsbruck.

## AUTHOR CONTRIBUTIONS

Conceptualization: V.L., S.S., and A.V. Methodology: S.S., N.K., I.R., J.H., M.S., M.E., J.S.R., F.F., and V.L. Investigation: S.S., N.K., I.R., P.Y.P., J.G.W., J.H., K.H. Visualization: S.S. Supervision: V.L., J.S.R., A.V., and F.F. Writing – original draft: S.S. and V.L.

## DATA AND MATERIALS AVAILABILITY

All data needed to evaluate the conclusions in the paper are present in the paper and/or the Supplementary Materials. Raw RNA-sequencing data will be deposited in the ArrayExpress collection in BioStudies under the accession number [to be provided upon publication]. All codes used in this study have been previously published and are referenced in the manuscript. Additional data related to this paper are available from the corresponding author upon request.

## Abbreviations

BCL2: B-cell lymphoma 2
BCR: B cell receptor
BIM: BCL2 interacting mediator of cell death
DEGs: differentially expressed genes
EdU: 5-ethynyl-2’-deoxyuridine
FACS: fluorescence-activated cell sorting
IgD: Immunoglobulin D
Ighv: Immunoglobulin heavy chain variable regiou
IgM: Immunoglobulin M
LOF: loss-of-function
MCL1: myeloid cell leukemia 1
MFI: mean fluorescence intensity
pAKT: phosphorylated protein kinase B
pH3: phosphohistone H3
TCF3/E2A: transcription factor 3 (E2A immunoglobulin enhancer-binding factors E12/E47)
TET2: ten-eleven translocation 2
VH: Variable Heavy chain
γH2AX: gamma-histone variant H2AX

