## Supplementary Figures for "TET2 loss promotes premalignant survival and clonal selection in MYC-driven B cell lymphoma"

### Supplementary Figure 1

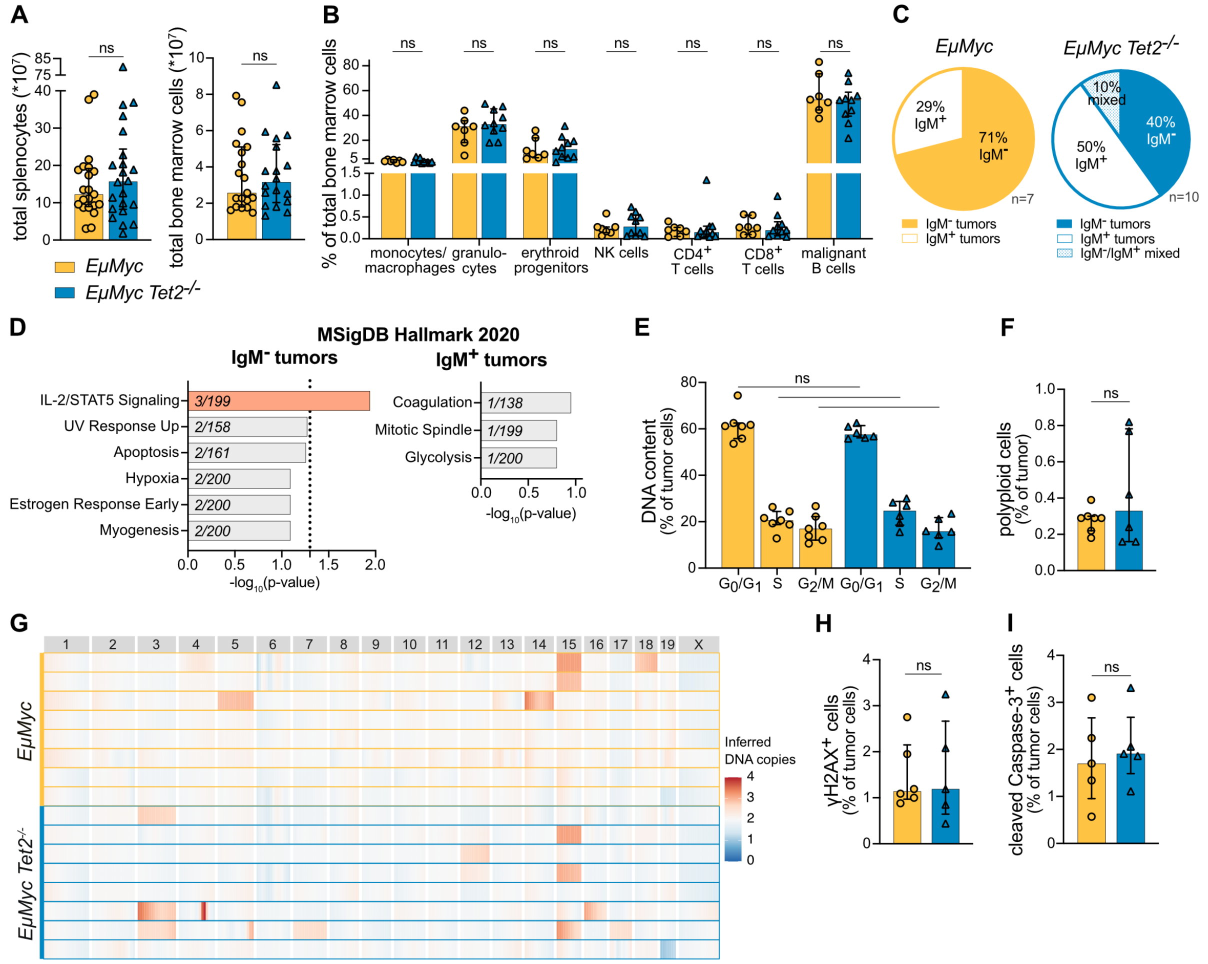


**Supplementary Figure 1: TET2 loss increases MYC-driven lymphoma penetrance and shifts the tumor spectrum toward IgM$^+$ cells. (A)** Absolute splenocyte counts (*10$^7$) (*EμMyc*: n=21, *EμMyc Tet2*$^{-/-}$: n=22) and total bone marrow cell counts (*10$^7$) (*EμMyc*: n=19, *EμMyc Tet2*$^{-/-}$: n=18) for each mouse. **(B)** Bone marrow immune cell composition assessed by flow cytometry using pooled cells of 2 femora and 2 tibiae: monocytes/macrophages (CD11b$^+$Gr1$^-$), granulocytes (CD11b$^+$Gr1$^+$), erythroid progenitors (nucleated Ter119$^+$), NK cells (TCRβ$^-$NK1.1$^+$), CD4$^+$ T cells (TCRβ$^+$CD4$^+$), CD8$^+$ T cells (TCRβ$^+$CD8$^+$), and malignant B cells (B220$^+$CD19$^+$) (*EμMyc*: n=7, *EμMyc Tet2*$^{-/-}$: n=10). **(C)** Lymphoma phenotype in the bone marrow. Mixed tumors were defined as tumors where neither IgM$^-$ nor IgM$^+$ cells constituted >80% of the total tumor population. **(D)** MSigDB Hallmark 2020 GO-term enrichment analysis (using Enrichr) for significantly DEGs (adjusted p-value<0.05 and absolute log$_2$(fold change)>1) from RNA-seq of splenic lymphoma cells comparing *EμMyc Tet2*$^{-/-}$ versus *EμMyc*, stratified by IgM$^-$ and IgM$^+$ tumors (*EμMyc*: n=3-5, *EμMyc Tet2*$^{-/-}$: n=4). The bar graph depicts the top enriched GO-terms ranked by adjusted p-value, with the number of overlapping DEGs contributing to each term indicated in italics inside each bar. **(E)** DNA content and **(F)** polyploid cell fraction of bone marrow cells, assessed by TO-PRO-3 staining via flow cytometry (*EμMyc*:

n=7 and *EμMyc Tet2*$^{-/-}$: n=6). **(G)** The aneuploidy score calculated from RNA-seq copy number data from splenic lymphoma cells, normalized to premalignant splenocytes (from Fig. 2E) of the corresponding genotype and IgM phenotype. Each horizontal line represents one individual tumor. **(H)** Flow cytometric analysis of DNA double strand breaks by γH2AX (*EμMyc*: n=6, *EμMyc Tet2*$^{-/-}$: n=5) and **(I)** apoptosis by cleaved Caspase-3 (*EμMyc*: n=5, *EμMyc Tet2*$^{-/-}$: n=5) in bone marrow cells. Bar graphs show median with interquartile range. Statistical significance was determined using (A, B) Mann-Whitney test, or (E, F, H, I) unpaired t-test, with Holm-Šidák correction for multiple comparisons. Normality was assessed using the Shapiro-Wilk test. ns = not significant.

### Supplementary Figure 2

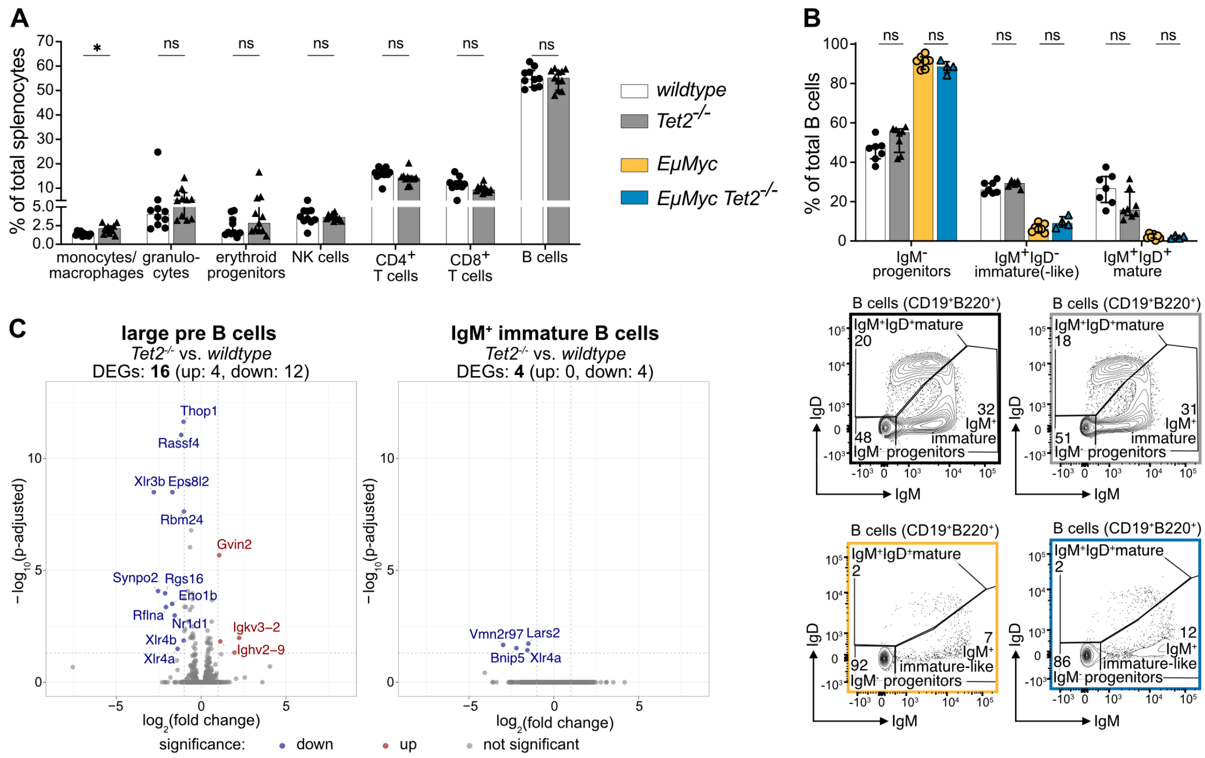

**Supplementary Figure 2: *Tet2* loss has limited effects in the absence of oncogenic MYC. (A)** Splenic immune cell composition assessed by flow cytometry: monocytes/macrophages ($CD11b^{+}Gr1^{-}$), granulocytes ($CD11b^{+}Gr1^{+}$), erythroid progenitors (nucleated $Ter119^{+}$), NK cells ($TCR\beta^{-}NK1.1^{+}$), $CD4^{+}$ T cells ($TCR\beta^{+}CD4^{+}$), $CD8^{+}$ T cells ($TCR\beta^{+}CD8^{+}$), and B cells ($B220^{+}CD19^{+}$) (*wildtype*: n=10, *Tet2*$^{-/-}$: n=11). **(B)** Bone marrow B cell subsets were assessed via flow cytometry: $IgM^{-}$ progenitors ($B220^{+}CD19^{+}IgM^{-}IgD^{-}$), $IgM^{+}IgD^{-}$ immature-(like) ($B220^{+}CD19^{+}IgM^{+}IgD^{-}$), and $IgM^{+}IgD^{+}$ mature ($B220^{+}CD19^{+}IgM^{+}IgD^{+}$) B cells. The upper graph summarizes all data (*wildtype*: n=7, *Tet2*$^{-/-}$: n=8, *EμMyc*: n=7, *EμMyc Tet2*$^{-/-}$: n=4), the lower panel shows representative dot blots for the IgM/IgD gate. **(C)** Volcano plots display RNA-seq derived transcriptional profiles of FACS-sorted splenic large pre B cells (left; $B220^{+}CD19^{+}IgM^{-}IgD^{-}cKit^{-}CD25^{+}$FSC-A) and $IgM^{+}$ immature (right; $B220^{+}CD19^{+}IgM^{+}IgD^{-}$) B cells. Comparisons between young (50 days old) *Tet2*$^{-/-}$ (n=3) and *wildtype* (n=3) mice were performed separately for each cell type. Axis ranges were kept identical across volcano plots to allow direct comparison. Significance was defined as adjusted p-value<0.05 and absolute $\log_2$(fold change)>1. Downregulated genes in *Tet2*$^{-/-}$ subsets are shown in blue; upregulated genes are shown in red. Bar plots show median with interquartile range. Statistical significance was determined using (A) Mann-Whitney test and (B) two-way ANOVA, with Holm-Šidák correction for multiple comparisons. Normality was assessed using the Shapiro-Wilk test. ns = not significant, *p<0.05.

**Supplementary Figure 3**

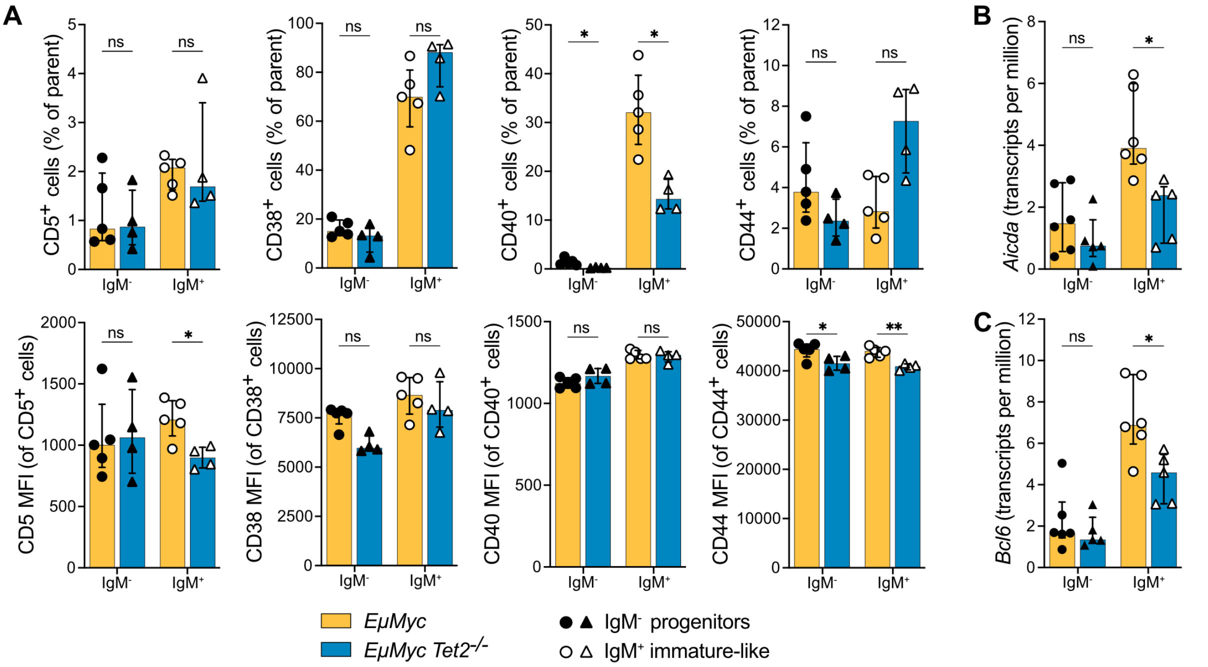


**Supplementary Figure 3: TET2 loss undermines IgM+ immature-like B cells maturation circuitry in *EµMyc* mice. (A)** Flow cytometric validation of selected B-lineage regulatory/maturation gene panel in IgM- progenitors (filled symbols; B220+CD19+IgM-IgD-) and IgM+ immature-like B cells (empty symbols; B220+CD19+IgM+IgD-). The upper panel displays the percentage of B cell subsets expressing the indicated markers, while the lower panel shows the geometric MFI within the respective marker-positive gates. Transcripts per million of **(B)** *Aicda* and **(C)** *Bcl6* are plotted, based on RNA-seq data from FACS-sorted IgM- progenitors and IgM+ immature-like B cells from the comparison of premalignant *EµMyc Tet2$^{-/-}$* (n=5) versus *EµMyc* (n=6) mice. Bar plots show median with interquartile range. Statistical significance was determined using unpaired t-test for (A) %CD40+, %CD44+, and MFI CD44, %CD5+, and MFI CD5, %CD38+ and (B, C), or Mann-Whitney test for (A) CD40 MFI and CD38 MFI, with Holm-Šidák correction for multiple comparisons. Normality was assessed using the Shapiro-Wilk test. MFI = mean fluorescence intensity, ns = not significant, *p<0.05, **p<0.005.

**Supplementary Figure 4**

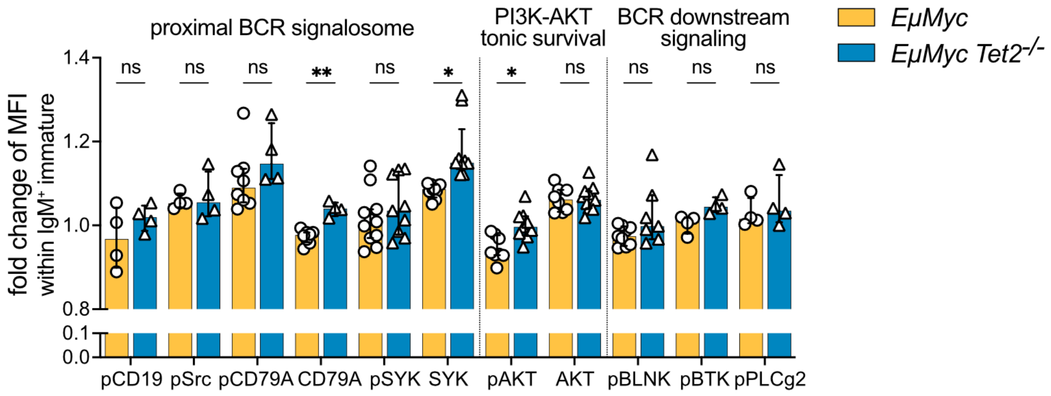


**Supplementary Figure 4: BCR-associated signaling in IgM$^+$ immature-like B cells.** Intracellular flow cytometric analysis of B cell receptor (BCR) signaling proteins in IgM$^+$ immature-like B cells (B220$^+$CD19$^+$IgM$^+$IgD$^-$) from *EμMyc* (n=4) and *EμMyc Tet2*$^{-/-}$ (n=4) mice. Data represent MFI normalized to the total B cell (CD19$^+$B220$^+$) gate. Bar plot shows median with interquartile range. Statistical significance was assessed using multiple unpaired t-test. Normality was evaluated using the Shapiro-Wilk test. ns = not significant, *p<0.05, **p<0.005.

### Supplementary Figure 5

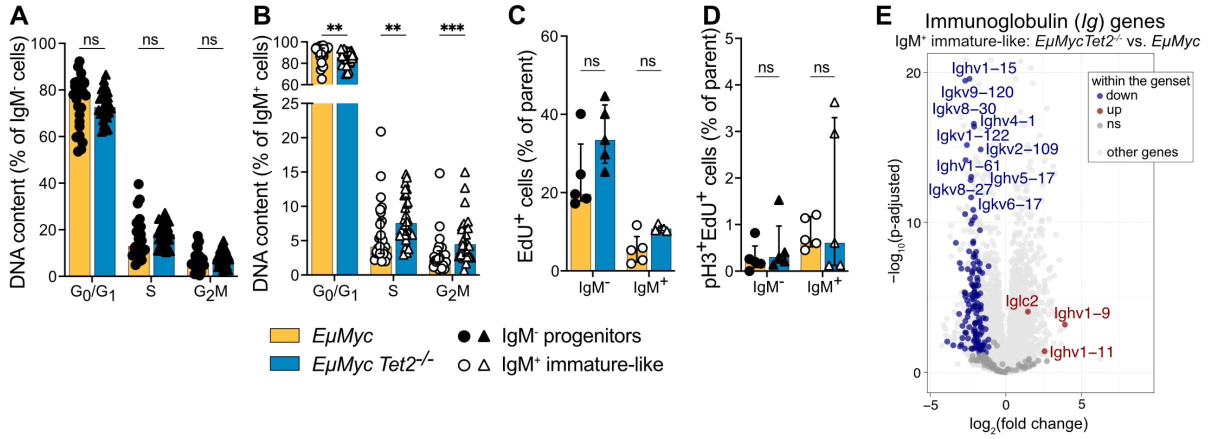


**Supplementary Figure 5: *Tet2* loss alters cell cycle kinetics within the IgM$^+$ immature-like B cell compartment.** Flow cytometric analysis of DNA content (cell cycle distribution) using TO-PRO-3 staining in **(A)** IgM$^-$ progenitors (B220$^+$CD19$^+$IgM$^-$IgD$^-$) and **(B)** IgM$^+$ immature-like B (B220$^+$CD19$^+$IgM$^+$IgD$^-$) cells from *EμMyc* (n=26) and *EμMyc Tet2*$^{-/-}$ (n=28) mice. Quantification of **(C)** EdU incorporation and **(D)** EdU incorporation combined with pH3$^+$ staining in IgM$^-$ progenitors and IgM$^+$ immature-like B cells. Total splenocytes from *EμMyc* (n=5) and *EμMyc Tet2*$^{-/-}$ (n=5) mice were labeled *in vitro* with EdU for 2 hours before analysis. **(E)** Volcano plot comparing transcriptional profiles, with a specific focus on immunoglobulin (*Ig*) genes. Volcano plot is based on the RNA-seq analysis of FACS-sorted IgM$^+$ immature-like (B220$^+$CD19$^+$IgM$^+$IgD$^-$) B cells comparing premalignant *EμMyc Tet2*$^{-/-}$ (n=5) versus *EμMyc* (n=6). Significant differential expression was defined as adjusted p<0.05 and absolute log$_2$(fold change)>1. Downregulated genes in *EμMyc Tet2*$^{-/-}$ subsets are shown in blue; upregulated genes are shown in red; non-*Ig* genes are depicted in dark grey, and all other genes in bright grey. For improved visualization, the volcano plot was cropped to highlight the most strongly deregulated *Ig* genes. Statistical significance was determined in all graphs using multiple Mann-Whitney tests, with Holm-Šidák correction for multiple comparisons. Normality was assessed using the Shapiro-Wilk test. ns = not significant, *p<0.05, **p<0.005, ***p<0.0005.
